# Improved imaging and preservation of lysosome dynamics using silver nanoparticle-enhanced fluorescence

**DOI:** 10.1101/2022.04.26.489585

**Authors:** Sumaiya A. Soha, Araniy Santhireswaran, Saaimatul Huq, Jayde Casimir-Powell, Nicala Jenkins, Gregory K. Hodgson, Michael Sugiyama, Costin N. Antonescu, Stefania Impellizzeri, Roberto J. Botelho

## Abstract

The dynamics of living cells can be studied by live-cell fluorescence microscopy. However, this requires the use of excessive light energy to obtain good signal-to-noise ratio, which can then photobleach fluorochromes, and more worrisomely, lead to photo-toxicity. Upon light excitation, noble metal nanoparticles such as silver nanoparticles (AgNP) generate plasmons, which can then amplify excitation in direct proximity of the nanoparticle’s surface and couple to the oscillating dipole of nearby radiating fluorophores, modifying their rate of emission and thus, enhancing their fluorescence. Here, we show that AgNP fed to cells to accumulate within lysosomes enhanced the fluorescence of lysosome-targeted Alexa488-conjugated dextran, BODIPY-cholesterol, and DQ-BSA. Moreover, AgNP increased the fluorescence of GFP fused to the cytosolic tail of LAMP1, showing that metal enhanced fluorescence can occur across the lysosomal membrane. The inclusion of AgNPs in lysosomes did not disturb lysosomal properties such as lysosomal pH, degradative capacity, autophagy and autophagic flux, and membrane integrity, though AgNP seemed to increase basal lysosome tubulation. Importantly, by using AgNP, we could track lysosome motility with reduced laser power without damaging and altering lysosome dynamics. Overall, AgNP-enhanced fluorescence may be a useful tool to study the dynamics of the endo-lysosomal pathway while minimizing photo-toxicity.

**eTOC:** Silver nanoparticles enhance fluorescence via surface plasmons. Here, we show that loading lysosomes with silver nanoparticles enhances the fluorescence of fluorochrome- and GFP-based molecular probes for lysosomes. This affords reduced excitation and exposure, diminishing photobleaching and phototoxicity, and preserving lysosome dynamics.

## Introduction

Live-cell fluorescence microscopy is an essential tool to determine and characterize the spatio- temporal dynamics of molecules, supra-molecular assemblies, organelles, and of entire cells (Lidke and Lidke, 2012; Jensen, 2013). Over the years, technological enhancements have pushed the boundary in the spatial resolution of cellular organization, the detection limits of cameras and photo-detectors allowing for single-molecule imaging, and image analysis that permits high-throughput, high-content and unbiased quantification (Lidke and Lidke, 2012; Boutros *et al*., 2015; Shashkova and Leake, 2017; Jing *et al*., 2021). These improvements depended and continue to depend on innovative manipulation of the physico-chemical properties of fluorochromes, hardware advances leading to superior camera sensitivity and detection, and a surge in computational power and better algorithms (Rust *et al*., 2006; Shtengel *et al*., 2009; Lidke and Lidke, 2012; Boutros *et al*., 2015; Peterson *et al*., 2016; Ouyang *et al*., 2018; Moen *et al*., 2019). Yet, challenges related to photo-bleaching and photo-toxicity remain when observing living samples by fluorescence microscopy, which lead to loss of signal, and more worrisomely, artifactual observations (Tinevez *et al*., 2012; Boudreau *et al*., 2016; Icha *et al*., 2017). These issues are most often caused by excessive light energy used during excitation, particularly blue light, which generates reactive oxygen species (ROS) that ultimately destroy fluorochromes and impair cell function (Tinevez *et al*., 2012; Waldchen *et al*., 2015; Boudreau *et al*., 2016; Icha *et al*., 2017). Thus, the need remains to develop brighter and more stable fluorochromes, more sensitive detection tools, and/or elegant manipulation of the physico-chemical properties of light and materials to further offset photo-bleaching and photo-damage.

Surface plasmons of noble metal nanoparticles are a promising means to enhance the emission intensity of fluorescent probes without the need to synthetically alter their chemical structure (Lakowicz *et al*., 2003; Bouhelier *et al*., 2004; Ekgasit *et al*., 2004b, 2004a; Petryayeva and Krull, 2011; Peterson *et al*., 2016; Hodgson *et al*., 2020). Plasmon excitation can be described as the collective oscillation of conduction band electrons on the surface of metal nanostructures following the incidence of a photon of appropriate wavelength (Barnes *et al*., 2003; Hao and Schatz, 2004; Liz-Marzán, 2006; Moores and Goettmann, 2006; Odom and Schatz, 2011). This effect translates into the enhancement of the electromagnetic field in proximity to the surface of the nanoparticles.

Silver nanoparticles (AgNP) are capable of interacting with light along the visible and near-infrared regions of the electromagnetic spectrum, and can significantly affect the photophysical processes underlying the emission of fluorescent molecules up to 200 nm, depending on the overlap of fluorophores’ emission spectra with the scattering component/s of the nanoparticles’ extinction spectra (Lakowicz *et al*., 2002, 2003, 2008; Aslan *et al*., 2005; Lakowicz, 2005; Anger *et al*., 2006; Fu and Lakowicz, 2007; Fu *et al*., 2007). This phenomenon is generally described as metal-enhanced fluorescence (MEF) and arises from near-field interactions between AgNP and a fluorochrome. This interaction ultimately enhances fluorescence by increasing the rate of fluorophore excitation (i.e., the enhanced electromagnetic field resulting from plasmonic excitation of AgNP increases the probability of excitation) and/or by increasing the radiative decay rate, which is proportional to the quantum yield of fluorescence (Hodgson *et al*., 2020). The latter mechanism occurs by coupling between the plasmon resonance oscillation of AgNP and the oscillating dipole of radiating fluorophores, effectively forming a radiative AgNP-fluorophore complex, often referred to as a “plasmophore”. Relative to the fluorochrome alone, the plasmophore exhibits a much lower excited state lifetime and a higher radiative decay rate, resulting in a higher quantum yield of emission (Lakowicz *et al*., 2008). Both components of MEF – increased excitation and increased quantum yield – ultimately contribute to enhanced brightness. Moreover, MEF can also improve photostability and reduce blinking (Malicka *et al*., 2003; Fu and Lakowicz, 2007).

Due to these reasons, the application of MEF by AgNP has considerable potential to improve fluorescence imaging of biological components. Due to the membrane impermeability of AgNP, MEF is best geared to enhance fluorescence of fluorochromes *in vitro*, or positioned on the cell surface, or within the endo-lysosomal compartments after pinocytosis. The endocytic pathway captures and processes extracellular and membrane-bound cargo and receptors for recycling or targeting to lysosomes (Huotari and Helenius, 2011; Naslavsky and Caplan, 2018; Norris and Grant, 2020). Therefore, AgNP are likely internalized by pinocytosis and trafficked to lysosomes. Lysosomes are a heterogeneous collection of acidic organelles that host numerous degradative enzymes, and are thus responsible for degrading internalized macromolecules, autophagic material, and particles engulfed by phagocytosis (Luzio *et al*., 2007; Kiselyov *et al*., 2012; Inpanathan and Botelho, 2019; Saffi and Botelho, 2019). Depending on the cell type, there are 100-1000 lysosomes per cell; these organelles are highly dynamic and can differ in their subcellular distribution, their motility, and fusion-fission cycling. These fusion and fission events can occur at the whole-organelle level or through membrane intermediates such as tubular extensions that aid in exchange of cargo (Bright *et al*., 2005; Balderhaar and Ungermann, 2013; Bissig *et al*., 2017; Choy *et al*., 2018).

Importantly, the labeling or illumination conditions typically required for fluorescence microscopy imaging have been shown to damage the endo-lysosomal system. For example, staining with acridine orange or Lysotracker, both used to detect acidic pH of lysosomes, can cause lysosome rupture when combined with light (Pierzyńska-Mach *et al*., 2014). Similarly, endo-lysosomal dynamics in macrophages was altered during live-cell spinning disc confocal imaging (Choy *et al*., 2018; Saffi *et al*., 2021). In part, this is mediated by reactive oxygen species formed during imaging, which can alter microtubule and actin dynamics (Choy *et al*., 2018; Saffi *et al*., 2021). In turn, altered microtubule and actin dynamics can disrupt lysosome fusion-fission cycles and transport (Bright *et al*., 2005; Johansson *et al*., 2007; Hong *et al*., 2015; Bissig *et al*., 2017; Saffi *et al*., 2021; Keren-Kaplan *et al*., 2022; Kumar *et al*., 2022). Fluorescence methods that reduce excitation energy and/or increase emission signal could reduce artifacts when investigating the endo-lysosomal system (Choy *et al*., 2018; Saffi *et al*., 2021).

Here, we postulated that AgNP would accumulate in lysosomes and could be used to enhance fluorescence of lysosome-targeted probes. In fact, we observed that AgNP enhanced the lysosomal fluorescence of several green-emitting dyes such as Alexa488-conjugated dextran, boron dipyrromethene (BODIPY)-labelled cholesterol, the dye-quenched DQ-BSA, which is used to measure protein degradation, and boosted the fluorescence of GFP fused to the cytosolic tail of LAMP1, a lysosomal membrane protein. As a result, we were able to reduce the excitation energy while obtaining good signal-to-noise ratio, allowing dynamic studies of lysosomes by preserving lysosome motility. Importantly, AgNPs did not seem to affect lysosomal properties such as pH, degradative capacity, membrane integrity, though an increased rate of tubulation of was observed in RAW cells. We propose that AgNP loading into lysosomes may be useful to study lysosome dynamics while mitigating photo-bleaching and photo-toxicity.

## Materials and Methods

### Synthesis of silver nanoparticles

Chemicals were purchased from Sigma-Aldrich and Fisher Scientific. To a clean, oven-dried 50 mL Erlenmeyer flask, 34.4 mL of MilliΩ H_2_O (Millipore Purification System), 4.0 mL of 10 mM trisodium citrate, 0.80 mL of 10 mM of 2-Hydroxy-4′-(2-hydroxyethoxy)-2- methylpropiophenone (Irgacure®-2959, I-2959) and 0.8 mL of 10 mM AgNO_3_ solutions were added. The mixture was degassed with N_2_ for 30 min, after which it was irradiated at 365 nm for 10 min in a Luzchem LCZ-4 photoreactor (14 bulbs, 100 W/m^2^) to yield a dark yellow solution of AgNP which was used as prepared. The obtained AgNP are ∼3 nm in diameter and are stable in solution up to 6 months (Stamplecoskie and Scaiano, 2010, 2012).

### Spectroscopy

Steady-state absorption spectra were recorded with an Agilent Cary 60 UV-visible spectrophotometer, using quartz cells with a path length of 1 cm. Steady-state emission and synchronous scattering spectra were recorded with an Agilent Cary Eclipse spectrofluorimeter in aerated solutions.

### AgNP casting and dye-embedded gelatin films

About 50-250 μL of AgNP suspension was dropcasted onto a clean, sterile 20x20 mm coverslip and dried overnight in the dark. We then prepared and filter-sterilized 0.2% (v/v) gelatin solution in ddH_2_O and added to a final concentration of 200 µM BODIPY^TM^ 493/503 (4,4- Difluoro-1,3,5,7,8-Pentamethyl-4-Bora-3a,4a-Diaza-s-Indacene; ThermoFisher) or 10 µg/mL BODIPY-cholesterol (ThermoFisher). We then added dropwise 150 μL of the BODIPY or BODIPY-cholesterol gelatin mixture onto coverslips previously coated with AgNP and incubated in the dark at room temperature for 5 min to solidify the gelatin. The coverslip was then lifted to pool the excess liquid at a corner and excess solution was aspirated by vacuum, leaving a thin film on the coverslip. The coverslips were left for an additional 20 min for the gelatin film to dry completely before imaging.

### Cell culture, plasmids, and transfection

The male-mouse macrophage-like RAW 264.7 cell line was obtained from ATCC (TIB-71, VA, USA) and cultured in DMEM with 5% heat inactivated fetal bovine serum, re-seeded every 2-3 days. The Cos7 cell line is a kidney fibroblast-like cell derived from a male-green monkey and was obtained from ATCC (CRL-1651) and were cultured in DMEM with 10% fetal bovine serum. Both cell types were plated at 30-50% confluency and used within 24-48 h. Plasmids were obtained from Addgene as follows: plasmid #34831 encoding LAMP1-GFP was a gift from Esteban Dell’Angelica, plasmid #73080 encoding galectin-3-GFP was a gift from Tamotsu Yoshimori, and plasmid #22418 encoding mCherry-GFP-LC3b was provided by Jayantha Debnath and previously characterized (Falcón-Pérez *et al*., 2005; N’Diaye *et al*., 2009; Maejima *et al*., 2013). Transfection was done with FuGENE (Promega, Madison, WI) as recommended by the manufacturer and cells were observed within 24 h.

### Pinocytosis of AgNP and labelling with lysosomal dyes

RAW and Cos7 cells were seeded at ∼70% confluency the day prior to imaging. Cells were then washed 1x with PBS and immersed with fresh 2 mL DMEM media supplemented with 5% FBS to which 50 or 250 μL of AgNP suspension was added. Cells were then incubated at 37 °C for 1h in the dark to permit internalization of AgNP suspension. After this, cells were washed 1x with PBS and immediately replenished with 1 mL of DMEM. Cells were then incubated with 10 µg/mL DQ-BSA (ThermoFisher) for 1 h, then washed in PBS and imaged in DMEM medium free of phenol red. Alternatively, cells were labelled with 10 µg/mL BODIPY-cholesterol for 40 min, then washed in PBS and imaged in DMEM medium free of phenol red. To label lysosomes with dextran conjugates, cells were all pulsed for 1 h and chased for 1 h with either with 100 µg/mL Alexa488-conjugated dextran (ThermoFisher) or 50 µg/mL Alexa546-dextran (ThermoFisher), unless otherwise stated below. We note here that we define lysosomes as organelles that labelled with fluid-phase markers and/or expression of LAMP1-GFP. We acknowledge that this likely encompasses a heterogeneous mixture of late endosomes, endolysosomes, and terminal lysosomes (Bright *et al*., 2005, 2005, 2016).

### Measuring relative lysosomal pH

RAW cells were seeded at 70% confluency the day prior to imaging. Cells were then washed 1x with PBS and immersed with fresh 2 mL DMEM supplemented with 5% FBS and 1% P/S, to which 250 μL of AgNP suspension was added. Cells were then incubated at 37L for 1 h in the dark to permit internalization of AgNP suspension. After this, cells were washed 1x with PBS and immediately replenished with 0.5 mL of DMEM. Cells were then co-labelled with 100 μg/mL pHRodo-Red-dextran and 100 μg/mL Alexa647-conjugated dextran for 1 h pulse and 30 min chase. Cells were then washed 3x with PBS and immediately replenished with 1 mL of DMEM. Cells were then treated with either 0.1% DMSO as vehicle control or 1 μM ConA for 15 min. Cells were imaged live in DMEM medium free of phenol red to quantify vacuolar pH changes, which were defined by measuring fluorescence intensity of dextran labelling.

### Induction of autophagy

RAW and Cos7 cells were seeded at 50-60% confluency two days prior to imaging. Cells were then transfected 24 h later with mCherry-GFP-LC3b. After 24 h of transfection, 250 μL of AgNP suspension were added and cells were then incubated at 37 L for 1 h in the dark to permit internalization of AgNP suspension. After this, cells were washed 1x with PBS and immediately replenished with 2 mL of DMEM. Cells were then left untreated, incubated for 3 h, and/or exposed to 1 μM torin1 for 3 h. Cells were imaged live in DMEM medium free of phenol red to quantify autophagic flux changes, which were defined by measuring autolysosome and autophagosome formation.

### Induction of lysosome damage

RAW 264.7 cells were transfected with galectin-3-GFP plasmid. Lysosomes were then labelled by incubating cells with 50 μg/mL Alexa647-conjugated dextran (ThermoFisher) for 1 h in complete media at 37°C in 5% CO2. Cells were washed PBS and resupplied with complete media for 30 min to chase the fluid-phase marker to lysosomes. Subsequently, cells were incubated with 50 μL or 250 μL of AgNP suspension and incubated at 37 °C for 1 h to permit internalization of AgNP suspension. After this, cells were washed 1x with PBS and immediately replenished with 1 mL of DMEM before live-cell imaging. As a positive control for lysosome damage, a subset of cells were treated with 1 mM Leu-Leu methyl ester hydrobromide (LLOMe; Sigma-Aldrich) for 2 h.

### Magic Red measurement of lysosome-degradative capacity

Raw cells were seeded to 70% confluency the day prior to imaging. On the day of imaging, cells were pulsed with 100 μg/mL of Alexa-647 dextran in fresh DMEM media, washed 3x with PBS, and chased in 2 mL fresh media for 30 min. Cells were then incubated in the dark with 0, 50, or 250 μL of AgNP in the dark at 37L for 1 h. After this, cells were washed 1x with PBS and were replenished with 480 μL fresh media. To detect cathepsin L activity, Magic Red (Bio-Rad) was reconstituted in DMSO to make a 260x stock concentration, followed by a dilution in sterile ddHlJO to make a 26x fold stock as per the manufacturer’s instruction. RAW cells were then incubated with Magic Red in a 1:26 ratio in media for 30 min. After this, cells were washed 1x with ice cold PBS and imaged live in phenol red-free DMEM media.

### LPS-induced lysosome tubulation

RAW264.7 cells were seeded at 70% confluency the day prior to imaging in DMEM supplemented with 5% FBS for 24 h at 37 UC in 5% CO_2_. Lysosomes were labelled by incubating cells with 50 μg/mL Alexa546-conjugated dextran (ThermoFisher) for 1 h in complete media at 37 UC in 5% CO_2_. Cells washed with PBS and resupplied with complete media for 30 min to chase the fluid-phase marker to lysosomes. Cells were then washed 1x with PBS and immersed with fresh 1 mL complete media to which 50 μL or 250 μL of AgNP suspension was added. Cells were then incubated at 37°C for 1 h in the dark to permit internalization of AgNP suspension. After this, cells were washed 1x with PBS and immediately replenished with 1 mL of DMEM. Cells were then incubated with 500 ng/ml lipopolysaccharide (LPS) from *Escherichia coli* (InvivoGen) for 2 h to induce lysosome tubulation and then washed in PBS and imaged in DMEM medium live.

### Live-cell spinning disc confocal microscopy

Cells were imaged by live-cell spinning disc confocal microscopy using a Quorum Diskovery Spinning Disc Confocal microscope system equipped with a Leica DMi8 microscope connected to a iXon Ultra 897 EMCCD BV camera and controlled by Quorum WaveFX powered by MetaMorph software (Quorum Technologies, Guelph, ON). Imaging was done with a 63x oil immersion objective and 100 µm pinhole disc. Imaging settings during each session were kept the same with typical settings as follows: electron-multiplying gain of 50, exposure of 100 ms, and the laser power set to 50 mW. Lower laser light exposure was used at 10 ms exposure, 10 mW laser power, and an electron-multiplying gain of 50. Single plane images were acquired with the brightfield channels and appropriate laser and filter settings. Images were 16-bit and 512 x 512.

### Incucyte imaging and quantification of confluency and CellTox Green staining

RAW macrophages were seeded into a 6-well plate at 30-50% confluency. The next day, cells were labelled with vehicle, 50 µL of AgNPs, or 250 µL of AgNPs as before, followed by 1 μL/mL CellTox Green (Promega). Alternatively, cells were treated with 5 µM staurosporine (Abcam, Cambridge, MA) and CellTox Green as a positive control for cell death. Immediately after adding the treatments and CellTox Green, cells were imaged using the Incucyte SX5 Live-Cell Analysis Instrument (Sartorius, Gottingen, Germany). Cells were imaged using the phase and green channels for 24 h every 30 min, using the 10x objective lens. Images were analysed at each frame by calculating the percent area covered by cells (phase imaging) and the number of CellTox Green-labelled cells using particle detection module after thresholding and subtracting background fluorescence.

### Image analysis and quantification

Image analysis was performed with ImageJ software with FIJI modules (National Institutes of Health, Bethesda, MD; (Schneider *et al*., 2012). The fluorescence intensity of 16-bit images or regions of interest (single cells) was measured by subtracting background. Due to variation between experiments in the absolute values in fluorescence, means were normalized as indicated. For visualization, images were window balanced with conditions that yielded highest intensity and propagated to other open images without altering pixel values. The number of images or cells quantified per condition and per experiment are indicated in the relevant figure legends.

### Lysosome positioning, particle tracking, and lysosome motility

Lysosome positioning was quantified in RAW macrophages expressing LAMP1-GFP. The cells were seeded at 75 % confluency the day prior to imaging. On the day of imaging the cells were treated with 50 µL or 250 µL AgNP for 1 h and imaged with spinning disc microscopy to acquire Z-stacks. The Z-stacks were collapsed in ImageJ/FIJI (NIH, Bethesda, MD). Single cells were outlined to measure the fluorescence intensity of the entire cell (shell 1). The regions of interest for each cell were then reduced by 10 pixels in three consecutive steps to form shells 2, 3 and 4, respectively. We then calculated the ratio of percent signal in shell 1 to shell 4.

Lysosome motility was quantified using TrackMate plug-in for ImageJ/FIJI. Briefly, individual cells were selected and subject to TrackMate analysis (Tinevez *et al*., 2017). Particle size was set to 10 pixels across all conditions, while thresholding was modified to track 5-10 puncta per cell, eliminating noise. The Simple LAP Tracker was selected and the linking max distance, gap-closing max distance, and gap closing max frame gap were maintained at the default values of 15, 15, and 2, respectively. No additional filters were applied to avoid bias. Velocity analysis was extracted and used to quantify lysosome motility.

### Statistical analysis

Experiments were repeated a minimum of three independent times or as indicated. For single-cell analysis, 15-50 cells were quantified per experiment per condition as indicated in figure legends. Unless otherwise indicated, means or normalized means and the associated standard error of the mean (SEM) of the independent experiments were calculated and statistically analysed. Unless otherwise stated, data was assumed to be normally distributed, Student’s t-test was used to compare experiments with two conditions, and one-way ANOVA and Tukey’s post-hoc test was used to test differences in means between three or more conditions. Statistical analysis and data presentation was done with GraphPad Prism version 9 and following suggestions by Prism based on the provided data.

## Results

### AgNP enhances BODIPY fluorescence in vitro

To assess the ability of AgNP to enhance fluorescence, we synthesized small AgNP (Ag “seeds”) averaging 3 nm in diameter through a photochemical method. Upon ultraviolet (365 nm) illumination, the free radical initiator I-2959 can undergo a Norrish II cleavage to produce ketyl radicals (Fig. 1A). To maximize the photochemical cleavage of I-2959, as well as minimizing quenching of the photoproducts by oxygen, the solutions were purged with N_2_ for 30 min prior to irradiation. The photogenerated ketyl radicals can then reduce Ag^+^ to Ag^0^, leading to the formation of AgNP.

**Figure 1:**
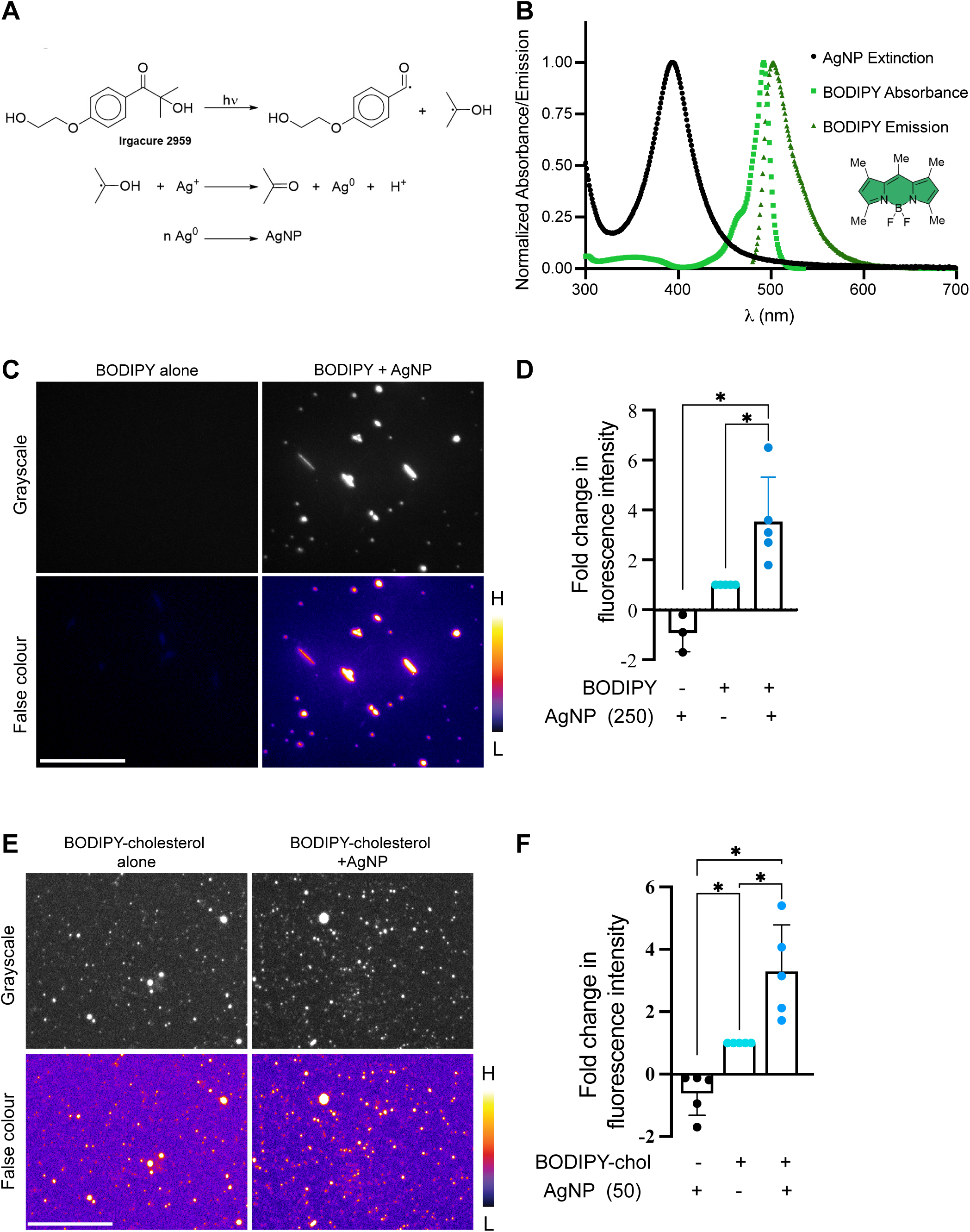
AgNP enhances fluorescence intensity of BODIPY embedded in a gelatin matrix. **A.** Schematic of the synthesis pathway for AgNP seeds. **B.** Normalized steady-state absorption (20 °C, CH_3_CN) and emission spectra of BODIPY^493/503^ (20 °C, CH_3_CN, λ_Ex_ = 460 nm), and extinction spectrum of AgNP (20 °C, water). **C, E.** A mixture of gelatin and BODIPY (C) or of gelatin and BODIPY-cholesterol (E) was casted on top of coverslips with no AgNP or with a AgNP-film dried onto the coverslip. Gelatin films were then imaged by spinning disc confocal microscopy. Images are shown as false-colour, where white-yellow is highest intensity and black-blue is lowest intensity. Scale bar = 10 µm. **D, F.** The mean fluorescence intensity of images was quantified including images with AgNP dried but no fluorochome added (first column). The means ± SEM from N=5 experiments are shown. The mean was normalized to the fluorochome-only condition. One-way ANOVA and Tukey’s post-hoc test were used to compare means, * p<0.05.

The extinction spectrum of AgNP (Fig. 1B, black trace) shows the absorption of the plasmon resonance band at 390 nm, while the nanoparticles are not fluorescent. We then sought to determine if these AgNP could enhance the fluorescence of model fluorochromes. For this purpose, we initially selected BODIPY, which displays maxima of absorption at ∼ 490 nm and emission at ∼ 520 nm (Fig. 1B, green traces). Previously, we demonstrated that the fluorescence of green-emitting probes based on the BODIPY molecular platform could experience MEF by large triangular silver nanoplates (∼ 80 nm), for which the surface plasmon band was centred at 660 nm (Dogantzis *et al*., 2020; Hodgson *et al*., 2020; Golian *et al*., 2021). In fact, large, non-spherical nanoparticles are known to exhibit increased scattering (Lakowicz, 2005; Lakowicz *et al*., 2008; Knoblauch *et al*., 2020). Nevertheless, while the scattering component of the extinction spectrum of AgNP tends to be greater for larger nanostructures of the same shape, it is the overlap between the emission spectrum of the dye and the nanoparticle scattering that is essential for MEF. In our case, the synchronous scattering (λ_Ex_ = λ_Em_) spectrum of AgNP (Dragan *et al*., 2013; Hodgson *et al*., 2020; Knoblauch *et al*., 2020) shows that the scattering component of the AgNP extinction overlaps well with the emission of BODIPY and, by extension, other dyes emitting at the similar wavelengths (Fig. S1, Supplementary Information).

In parallel, spectral overlap between the emission of the fluorophore and the absorption component of the nanoparticle’s extinction would be expected to quench fluorescence through the formation of non-radiative plasmophores. As shown in Fig. 1B however, minimal overlap exists between the fluorescence emission of BODIPY and the intrinsic absorption (where the synchronous scattering decreases) of AgNP in this system. It is worth noting here that the underlying requirement for MEF is ultimately the activation of surface plasmons, which can be achieved by the emission of nearby fluorophores, or by the same far-field irradiation used to excite the organic dye (in this work, λ_Ex_ = ∼488 nm). Although the extinction of AgNP at 488 nm is weak (Fig. 1B), we do not exclude a synergistic contribution of direct plasmon resonance excitation.

Next, we embedded BODIPY in a gelatin film, placing this onto a coat of AgNP dry casted onto a coverslip to determine if AgNP could boost the fluorescence of BODIPY fluorochromes *in vitro*. In this setup, the AgNP coating on the coverslips forms a separate layer onto which the BODIPY-containing gelatin film is formed, allowing MEF to occur in a spatially defined manner. First, we observed that gelatin films encasing the BODIPY dye and placed onto a coverslip without AgNP emitted low and diffused fluorescence by confocal imaging, but which was higher than coverslips containing only AgNP without fluorophore (Fig. 1C, 1D). In comparison, bright BODIPY spots were observed of different sizes and morphologies in the presence of AgNPs (Fig. 1C, 1D). In fact, the total fluorescence intensity of the image fields containing both BODIPY and AgNP was significantly higher (∼4x) than with BODIPY-alone, while both were higher than AgNP alone (Fig. 1C). To complement this experiment, we also used a similar strategy for BODIPY-cholesterol embedded in gelatin films placed on coverslips coated with AgNP and compared these to gelatin films with BODIPY-cholesterol on uncoated coverslips. We observed that BODIPY-cholesterol alone formed fluorescence puncta, possibly corresponding to cholesterol aggregates (Fig. 1E). This condition produced higher fluorescence than AgNP alone (Fig. 1F). By contrast, the fluorescence intensity of gelatin-fields containing BODIPY-cholesterol and placed on coverslips coated with AgNP was ∼3-fold higher than BODIPY-cholesterol alone (Fig. 1F). Overall, these data suggest that AgNPs enhance the fluorescence of free or conjugated BODIPY *in vitro*.

### AgNP are non-toxic to macrophages

We then tested if macrophages allowed to pinocytose AgNP would display signs of cytotoxicity before determining if AgNP could enhance fluorescence intracellularly. We used RAW macrophages as our primary cell model because of their proficiency for fluid phase endocytosis and trafficking to lysosomes. To test for cytotoxicity and based on an estimate of 1.261 x 10^11^ particles per µL suspension (see Supplementary Information and related discussion), we exposed cells to ∼6.3x10^12^ AgNP in 50 µL (6.3 penta-nanoparticles; P-np) or 3.2x10^13^ AgNP in 250 µL (32 P-np) to pinocytose over 24 h. During this period, cells were imaged using the Incucyte microscopy incubator system to measure cell growth and cell death. The former was tracked by change in confluency and the latter by enumerating CytoTox Green objects (non-viable cells). We observed no significant difference in the increase in cell confluency over 24 h between resting macrophages and those exposed to 50 or 250 µL of AgNP suspension (Fig. 2A, 2B, Videos 1, 2 and 3). Similarly, there was no apparent difference in the number of CytoTox Green objects that reflect non-viable cells between the three treatments (Fig. 2A, C, Videos 1, 2, and 3). In comparison, cells treated with staurosporine suffered a decline in confluency and were labelled with CytoTox Green in large numbers (Fig. 2, Video 4). Overall, these data intimate that treatment of cells with a suspension of AgNP was not significantly cytotoxic to RAW macrophages.

**Figure 2:**
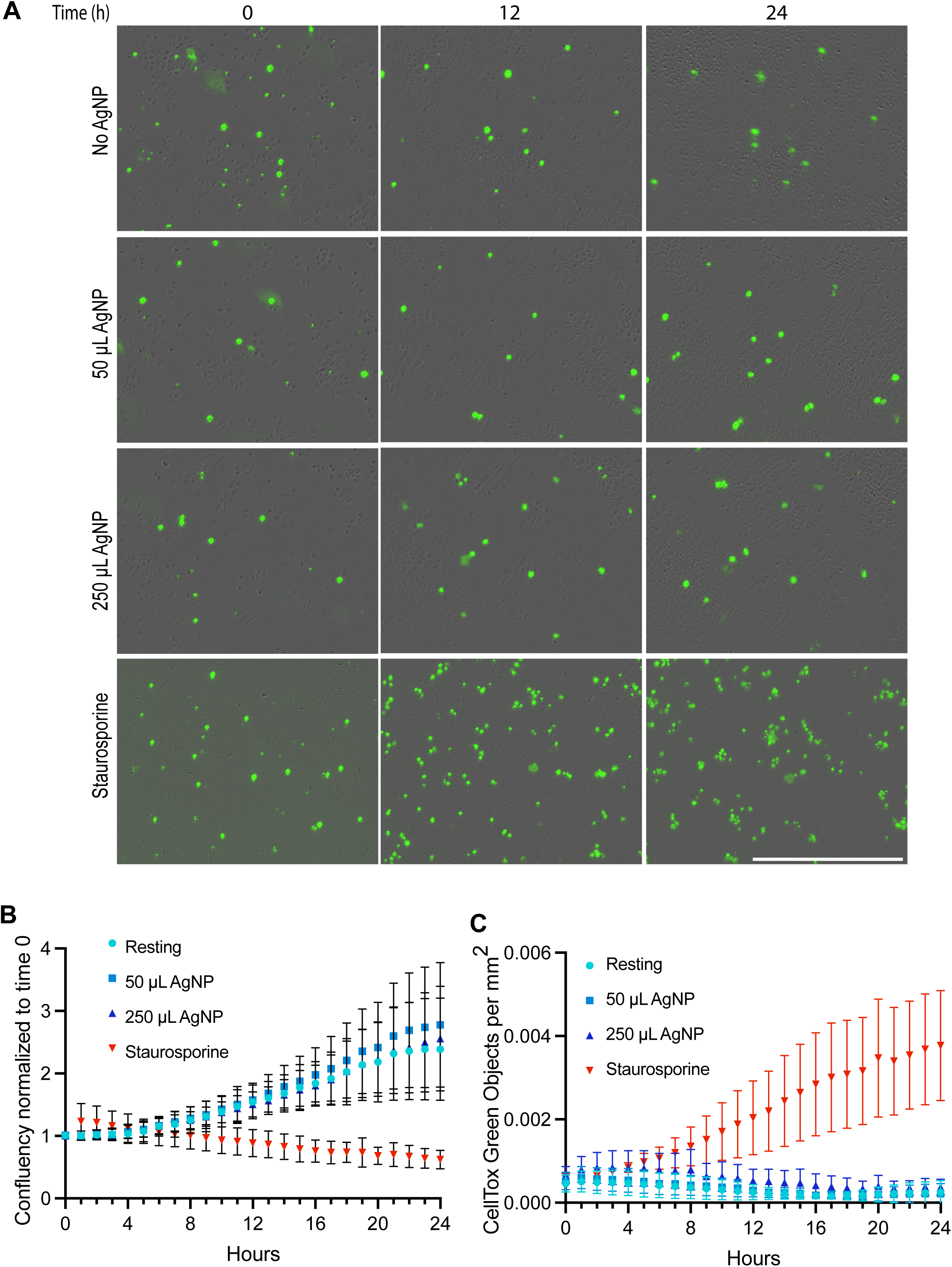
AgNP do not cause cytotoxicity in macrophages. **A.** RAW macrophages were kept resting, or they were treated over 24 h with 50 µL AgNP, 250 µL AgNP, or 5 µM staurosporine, while imaging in an Incucyte Incubator System. Confluency was tracked as an indicator of cell growth, while then number of CellTox Green particulates indicated cell death. Still frames at the indicated time points and treatments are shown. Scale bar = 400 µm. **B.** Percent confluency over 24 h normalized to resting cells. **C.** CellTox Green particles normalized to respective confluency for each condition shown. For B and C, shown are the mean ± SEM of N=4.

### AgNP enhance fluorescence of lysosome probes in macrophages

We then proceeded to test if AgNP accumulation in lysosomes could lead to metal-enhanced fluorescence in cells. To account for heterogeneity in uptake, accumulation of dye and AgNP, and possible interference due to proximity of AgNP to each other, we labelled macrophages with two different doses of AgNP: 50 µL or 250 µL of AgNP, as above. Subsequently, we labelled cells with either BODIPY-cholesterol, DQ-BSA, or Alexa488-conjugated dextran, followed by spinning disc confocal imaging, and analysis of the total fluorescence intensity per cell. First, cells were allowed to internalize 10 µg/mL BODIPY-cholesterol for 40 min and chased for 10 min to accumulate BODIPY-cholesterol on lysosomes (Hölttä-Vuori *et al*., 2016). Importantly, we observed an increase in fluorescence intensity of BODIPY-cholesterol in cells labelled with AgNPs (Fig. 3A, B). We do note that cholesterol is exported from lysosomes, creating a more diffused signal. Second, we labelled lysosomes with 10 µg/mL DQ-BSA for 1 h DQ-BSA, a molecular probe used to measure degradation within lysosomes that yields a fluorescent, green BODIPY once BSA is degraded in lysosomes (Marwaha and Sharma, 2017). Again, macrophages pre-loaded with AgNPs displayed higher DQ-BSA total fluorescence per cell relative to macrophages without AgNP in their lysosomes (Fig. 3C, D). Finally, to determine if AgNP could enhance a non-BODIPY dye in lysosomes with similar spectral properties, we incubated cells with 100 µg/mL Alexa488-conjugated dextran for 1 h and chased for 1h. As before, we observed higher average fluorescence of Alexa488-dextran in cells preincubated with AgNP relative to naïve cells, particularly those loaded with 250 µL AgNP (Fig. 3E, F). Overall, these data indicate that loading AgNP within lysosomes in macrophages can enhance fluorescence intensity, though we note that the dosage of AgNP that produces the best fluorescence enhancement may vary for each type of probe and should be optimized for each fluorescent compound.

**Figure 3:**
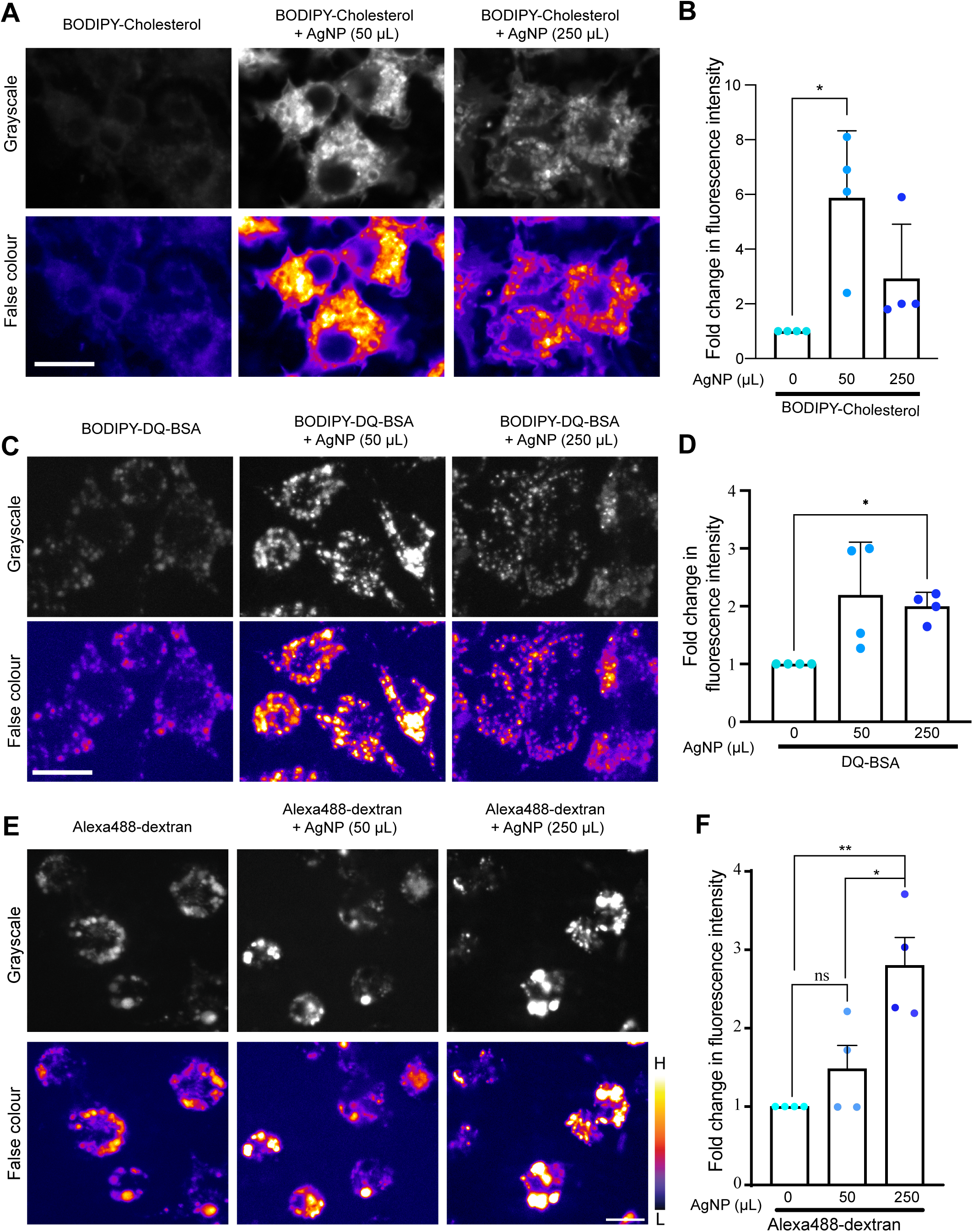
AgNP-enhanced fluorescence of BODIPY-cholesterol, DQ-BSA and Alexa488- conjugates in RAW macrophages. **A, C, E.** RAW cells were fed vehicle, 50 µL AgNP, or 250 µL AgNP, followed by 10 µg/mL BODIPY-cholesterol for 40 min and chased for 10 min (A), or 10 µg/mL DQ-BSA for 1 h (C), or 100 µg/mL Alexa488-conjugated dextran (E). Cells were then imaged by spinning disc confocal microscopy. Images are shown in grayscale (top) or false-colour (bottom), where white-yellow is highest intensity and black-blue is lowest intensity. Scale bar = 10 µm. **B, D, F.** Quantification of fluorescence intensity of BODIPY-cholesterol per cell (B), or DQ-BSA per cell (D), or Alexa488-dextran per cell (F), normalized to cells without AgNP. Shown is the mean ± SEM from N=4 experiments, where 15-30 cells were quantified per condition per experiment. One-way ANOVA and Tukey’s post-hoc test were used to compare means, * p<0.05, ** p<0.01.

### AgNP-enhanced fluorescence occurs across the lysosomal bilayer

Our observations suggest that AgNP can enhance fluorescence intensity of molecular probes within lysosomes. We next tested if AgNP could enable metal-enhanced fluorescence of GFP across the lysosomal membrane by expressing LAMP1-GFP, whereby the GFP is located on the cytosolic C-terminal domain of LAMP1 (Falcón-Pérez *et al*., 2005). Assuming GFP is 10 nm from the plane of the membrane and the lysosomal bilayer, inclusive of the glycocalyx is 30 nm thick, then GFP molecules should be within the effective distance for fluorescence enhancement by AgNP. To test this, we transfected RAW macrophages with plasmids encoding LAMP1-GFP, followed by exposing cells to vehicle, 50 µL or 250 µL AgNP seeds, as before. Cells were imaged and analysed by measuring the total LAMP1-GFP fluorescence using collapsed z-stacks. Notably, we observed a significant increase in the mean total LAMP1-GFP fluorescence in cells that accumulated AgNP relative to vehicle-only (Fig. 4A, B). To determine if metal enhanced fluorescence could also occur in other cell types, we transfected Cos7 cells with plasmid encoding LAMP1-GFP. As with RAW cells, we also observed a boost in LAMP1-GFP fluorescence in Cos7 cells containing AgNP (Fig. 4C, D). Altogether, our observations indicate that AgNP can enhance GFP fluorescence across the lysosomal bilayer. This is consistent with the emission spectrum of GFP with a maximum at λ_Em_ = 510 nm, which is not dissimilar from the fluorescence signature of BODIPY (Fig. 1B) and, thus, suitable to experience MEF by our AgNP.

**Figure 4:**
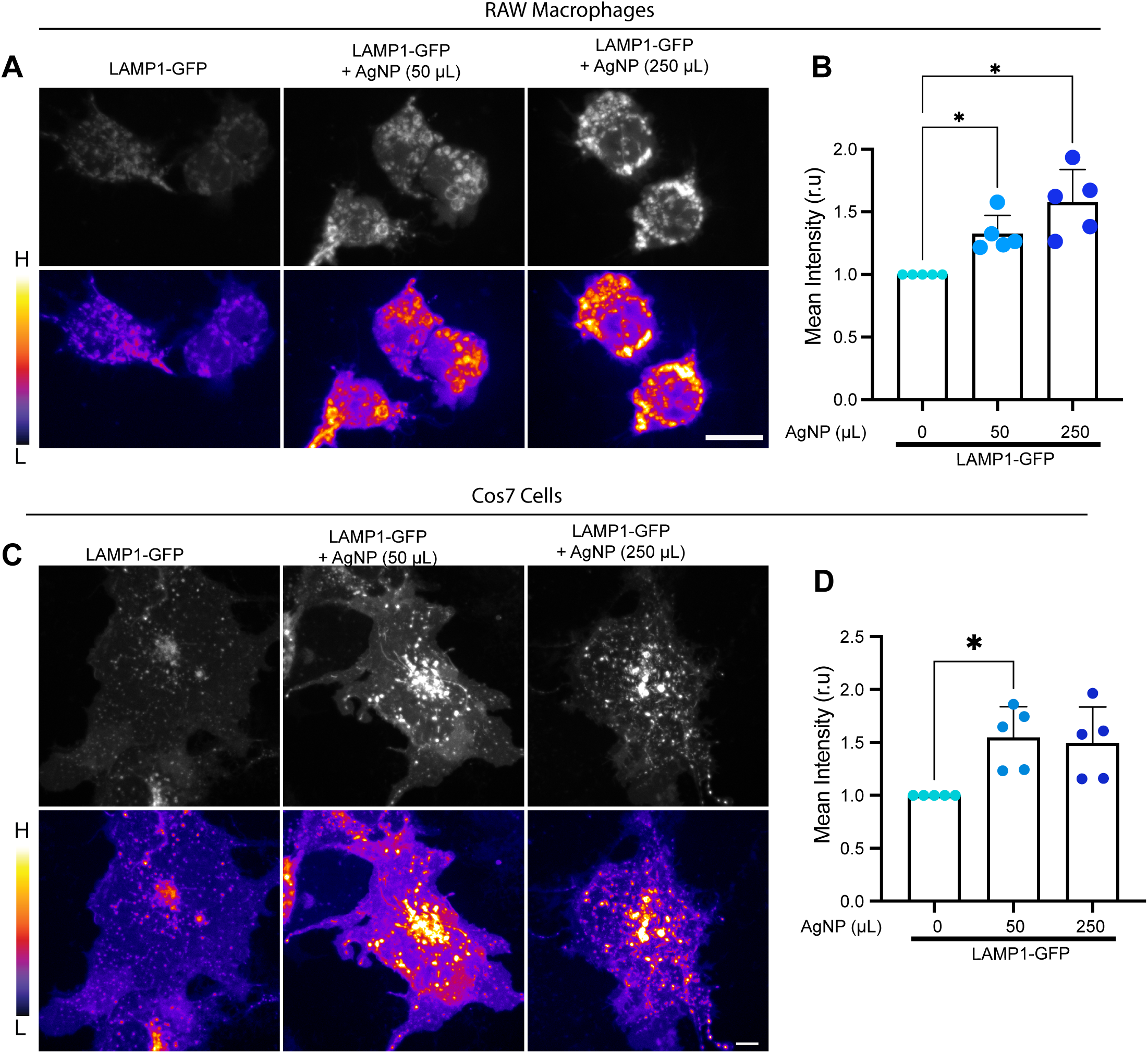
AgNP boosts fluorescence of LAMP1-GFP. **A, C.** RAW cells (A) and Cos7 cells (C) were transfected with plasmid encoding LAMP1-GFP. After 24 h, cells were loaded with vehicle, 50 µL AgNP, or 250 µL AgNP. Live-cells were then imaged by spinning disc confocal microscopy and are shown as grayscale (top row) or false-colour (bottom row) as before. Scale bar = 10 µm. **B, D**. The fluorescence intensity of LAMP1-GFP in RAW cells (B) or Cos7 cells (D) was quantified and normalized to control cells with no AgNP. Shown is the mean ± SEM from N=5 independent experiments, where 40-50 cells per condition per experiment were quantified. Means were then tested by one-way ANOVA, followed by Tukey’s post-hoc test, where * is p<0.05.

### AgNPs do not perturb lysosome properties

Our observations suggest that pinocytosis of AgNPs can enhance the fluorescence intensity of lysosome-targeted probes with spectral properties similar to BODIPY. We next systematically examined several lysosomal properties to determine the risk of artifacts caused by feeding cells AgNPs. First, using galectin-3-GFP as an indicator of gross lysosome damage (Chauhan *et al*., 2016), we did not observe evidence of lysosomal damage upon AgNP treatment as seen by the paucity of galectin-3-GFP puncta, while treatment with LLMeO, a known lysosome damaging agent, caused extensive accumulation of galectin-3-GFP puncta (Fig. 5).

**Figure 5:**
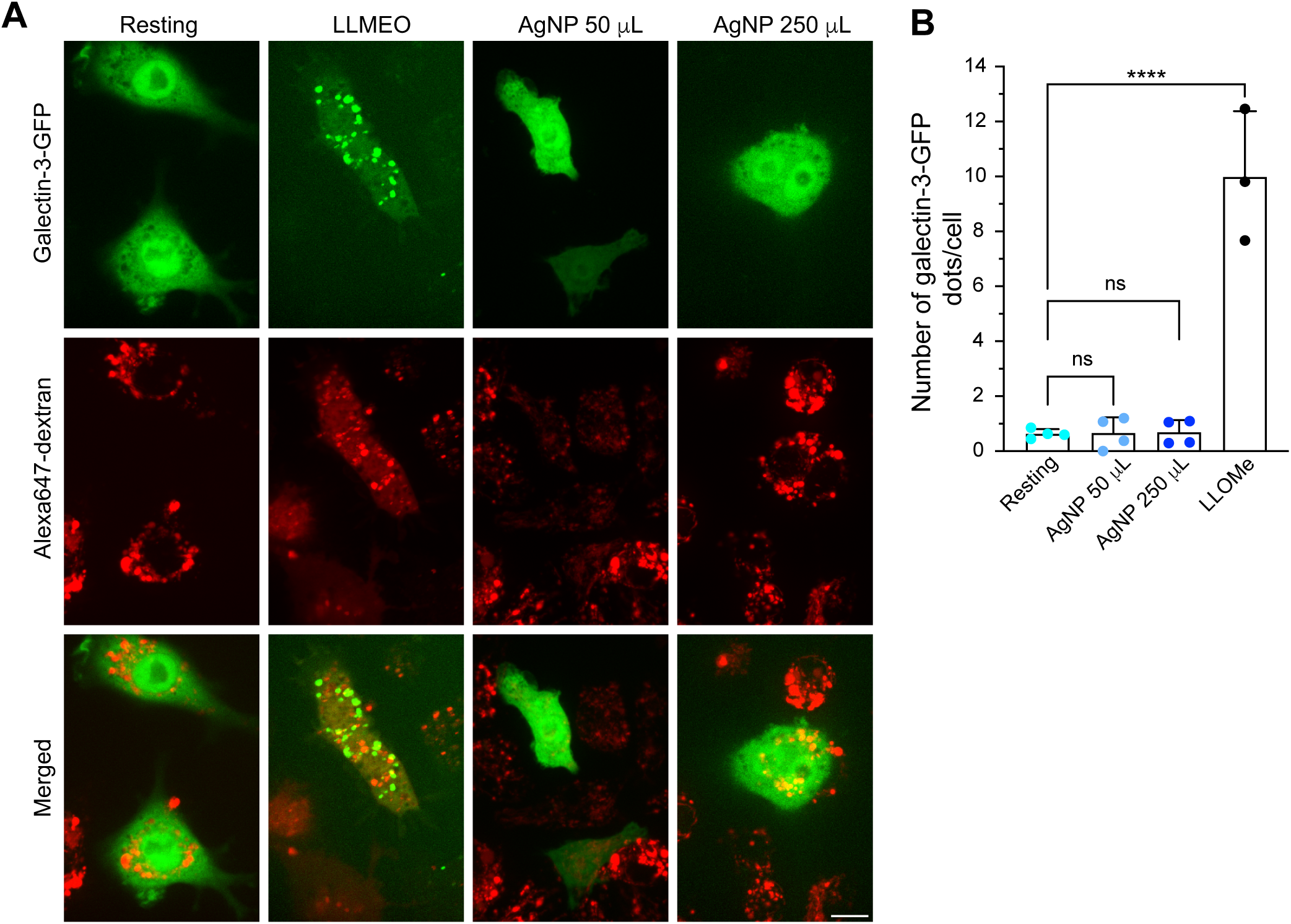
AgNPs do not induce lysosome damage in macrophages. RAW macrophages expressing galectin-3-GFP were labelled with 50 μg/mL Alexa647-conjugated dextran and then exposed to either 1 mM LLOMe for 2 h or with 50 μL or 250 μL AgNP for 1 h. Live-cell imaging was done by spinning disc confocal. Scale bar = 10 µm. **A.** Galectin-3-GFP is shown in green (top), Alexa647–dextran in red (middle) and the superimposition of the two fluorophores is shown as merge of the two channels (bottom). **B.** Mean number of galectin-3-GFP puncta per cell based on 25–30 cells per condition per experiment from N = 3 independent experiment (LLOMe) or N=4 for AgNP. One-way ANOVA followed by Tukey’s post-hoc test was used, where **** is p<0.001.

We then defined if more subtle disruption of lysosome properties occurred in cells loaded with AgNPs. To evaluate if AgNPs affected lysosomal pH, we labelled lysosomes of RAW cells with pHRodo Red- and Alexa647-conjugated dextrans with or without uptake of 250 µL of AgNPs. The ratio of pHRodo Red (pH sensitive) to Alexa647 (pH insensitive) was then used to estimate the relative lysosomal pH under these two conditions, finding no statistical difference between cells with and without AgNPs (Fig. 6). In comparison, addition of 1 µM concanamycin A to inhibit the V-ATPase alkalinized lysosomes (Fig. 6).

**Figure 6:**
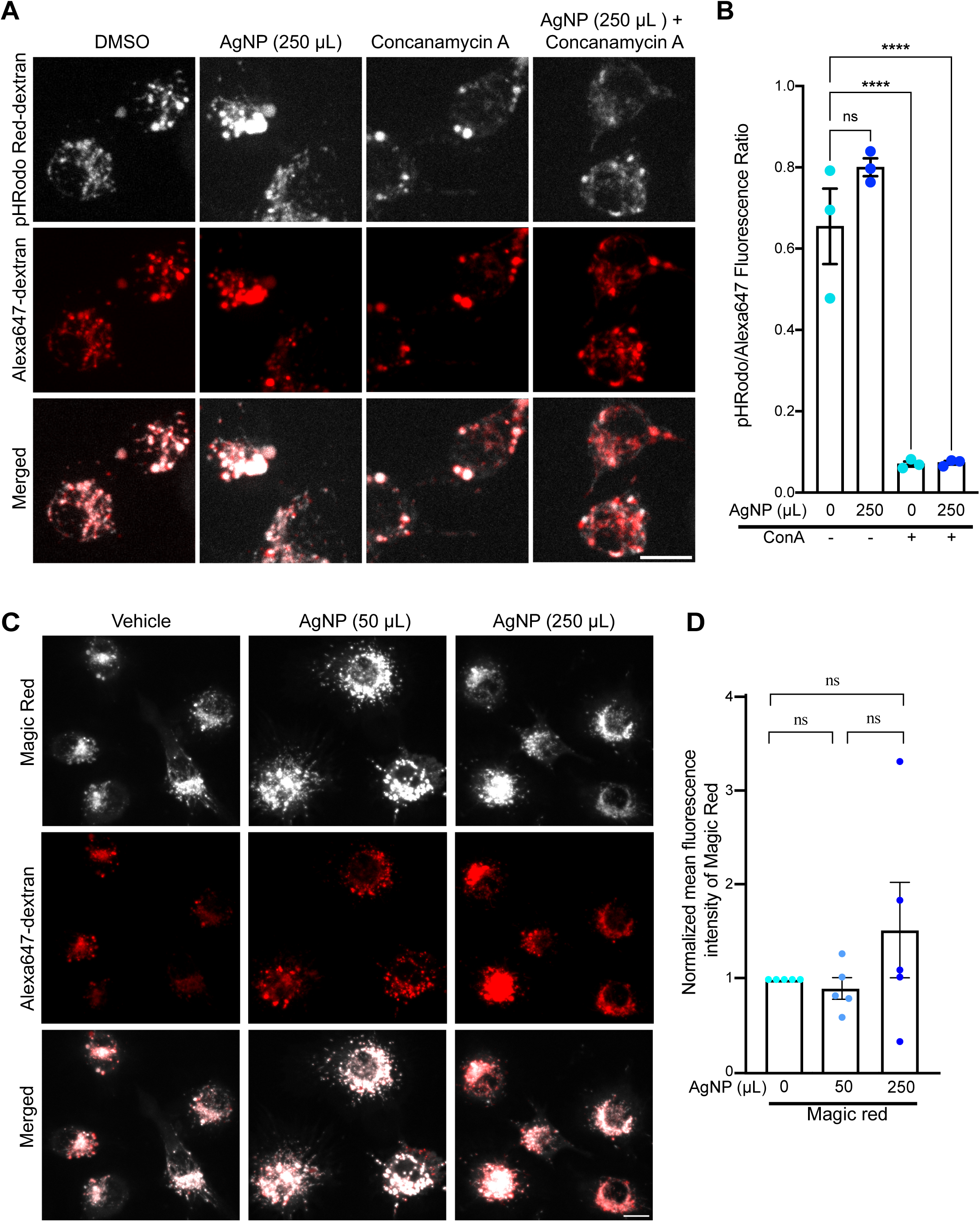
AgNP does not alter the pH and degradative properties of lysosomes in macrophages. **A.** RAW cells were fed vehicle or 250 µL AgNP for 1 h, followed by endocytosis and chase of 100 µg/mL Alexa647-dextran and 100 µg/mL pHRodo Red-dextran. Cells were then treated with vehicle or 1 µM Concanamycin A (ConA) and imaged using spinning disc confocal microscopy. pHRodo Red-dextran is shown in grayscale (top), Alexa647-dextran in red (middle), and the superimposition of the two fluorophores is shown as merge of the two channels (bottom). **B.** Ratio of pHRodo Red-dextran to Alexa647-dextran was quantified based on a minimum of 50 cells per condition per experiment. Shown is the mean ± SEM from N=3 independent experiments. One-way ANOVA and Tukey’s post-hoc test was used to compare means, **** p<0.001. **C.** RAW cells were treated with vehicle, or 50 µL AgNP, or 250 µL AgNP for 1 h, followed by endocytosis of 100 µg/mL Alexa647-dextran for 1 h and chased for 1 h, followed by Magic Red labelling, and imaged using spinning disc confocal microscopy. Magic Red is shown in grayscale (top), Alexa647-dextran in red (middle), and the superimposition of the two fluorophores is shown as merge of the two channels (bottom). For A and C: scale bar = 10 µm. **D.** Mean fluorescence of Magic Red was quantified and normalized to no AgNP condition, where a minimum of 25 cells per condition per experiment were quantified. Shown is the mean ± SEM from N=5 independent experiments. One-way ANOVA and Tukey’s post-hoc test were used to compare means, where no significant difference in pH was observed.

Following, we investigated the impact of AgNPs on lysosome degradative capacity using Magic Red Cathepsin L probe, which given its spectral properties is not expected to undergo AgNP-induced MEF. Importantly, there was no statistical difference in the average Magic Red fluorescence between cells loaded with or without AgNPs (Fig. 7). We then tested autophagy and autophagic flux by expressing mCherry-GFP-LC3b chimera (N’Diaye *et al*., 2009). The mCherry signal labels all autophagosomes, but because GFP fluorescence is quenched by low pH, one can differentiate between immature autophagosomes (mCherry^+^ GFP^+^ puncta) and autolysosomes (mCherry^+^ GFP^-^puncta) (Kimura *et al*., 2007). Nonetheless, loading AgNPs into RAW cells did not induce autophagy nor impact autophagic flux as measured by the number of LC3b-mCherry puncta (Fig. 8A, B) and the number of GFP+ mCherry+ puncta (Fig. 8A, C). We note that RAW cells had high-basal autophagy and that surprisingly torin1 treatment did not increase the number of autophagosomes (Fig. 8). To determine if this was a cell-specific issue, we tested autophagy and autophagic flux in Cos7 cells, which are also easier to transfect. Here, we observed that torin1 treatment induced autophagy (more LC3b-mCherry puncta upon torin1 incubation, Sup. Fig. S2). However, we once again did not observe a difference in autophagy or autophagic flux in Cos7 cells exposed to AgNPs (Supplemental Fig. S2).

**Figure 7:**
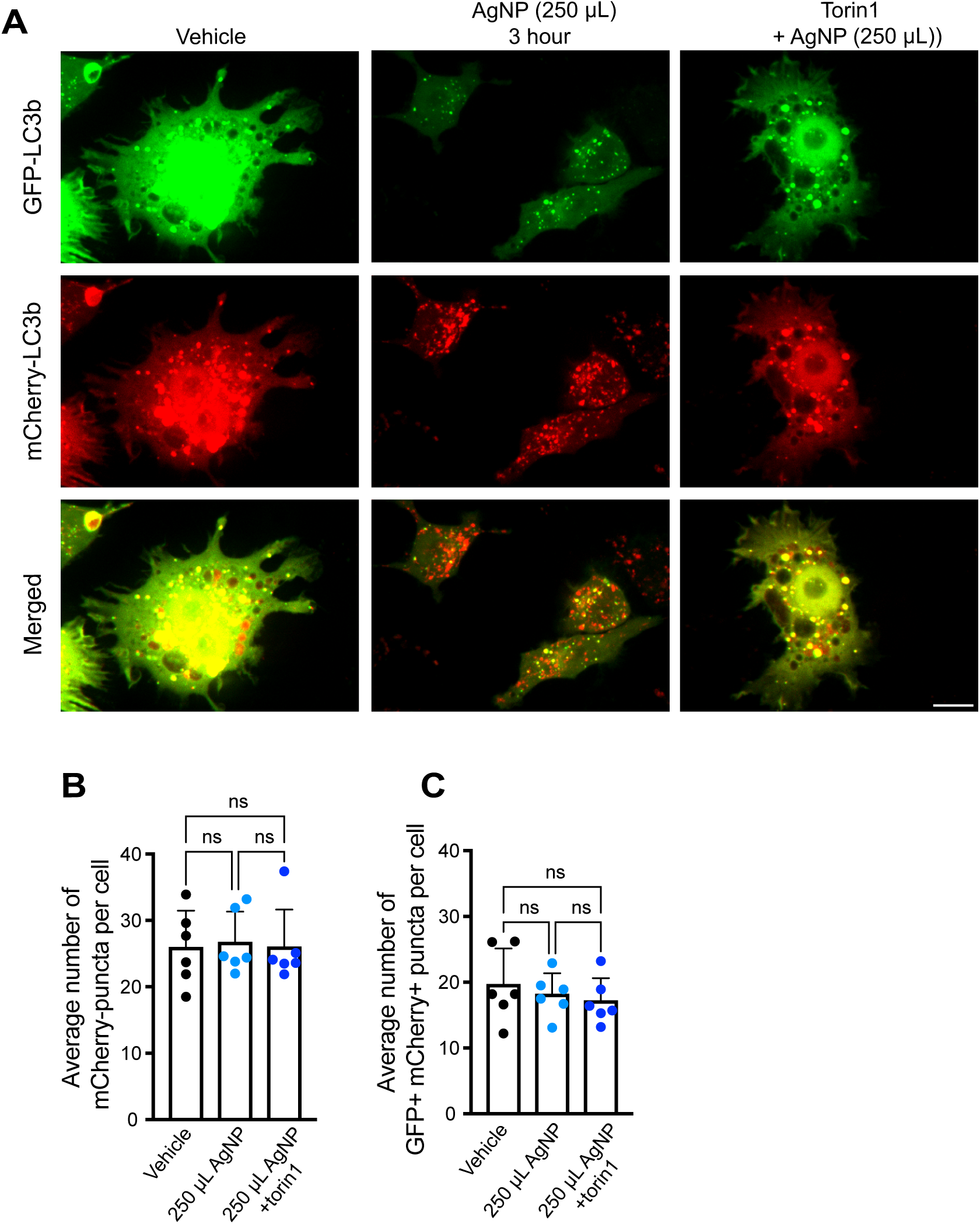
AgNP do not alter basal autophagy and autophagic flux in RAW cells. RAW cells were transfected with mCherry-GFP-LC3b plasmid and then exposed to vehicle, 250 µL AgNP, or 250 µL AgNP and 1 µM Torin1 for 3 h. Cells were then imaged live using spinning disc confocal microscopy. Scale bar = 10 µm. **A.** GFP signal is shown in green (top), while mCherry fluorescence is shown in red (middle), and the superimposition of the two fluorophores is displayed in the bottom. **B, C.** Mean number of mCherry puncta (B) and GFP+ mCherry+ puncta (C) were quantified from a minimum of 50 cells per condition per experiment. Shown is the mean ± SEM from N=6 independent experiments. One-way ANOVA and Tukey’s post-hoc test were used to compare means with no significant difference in autophagy and autophagic flux observed between.

**Figure 8:**
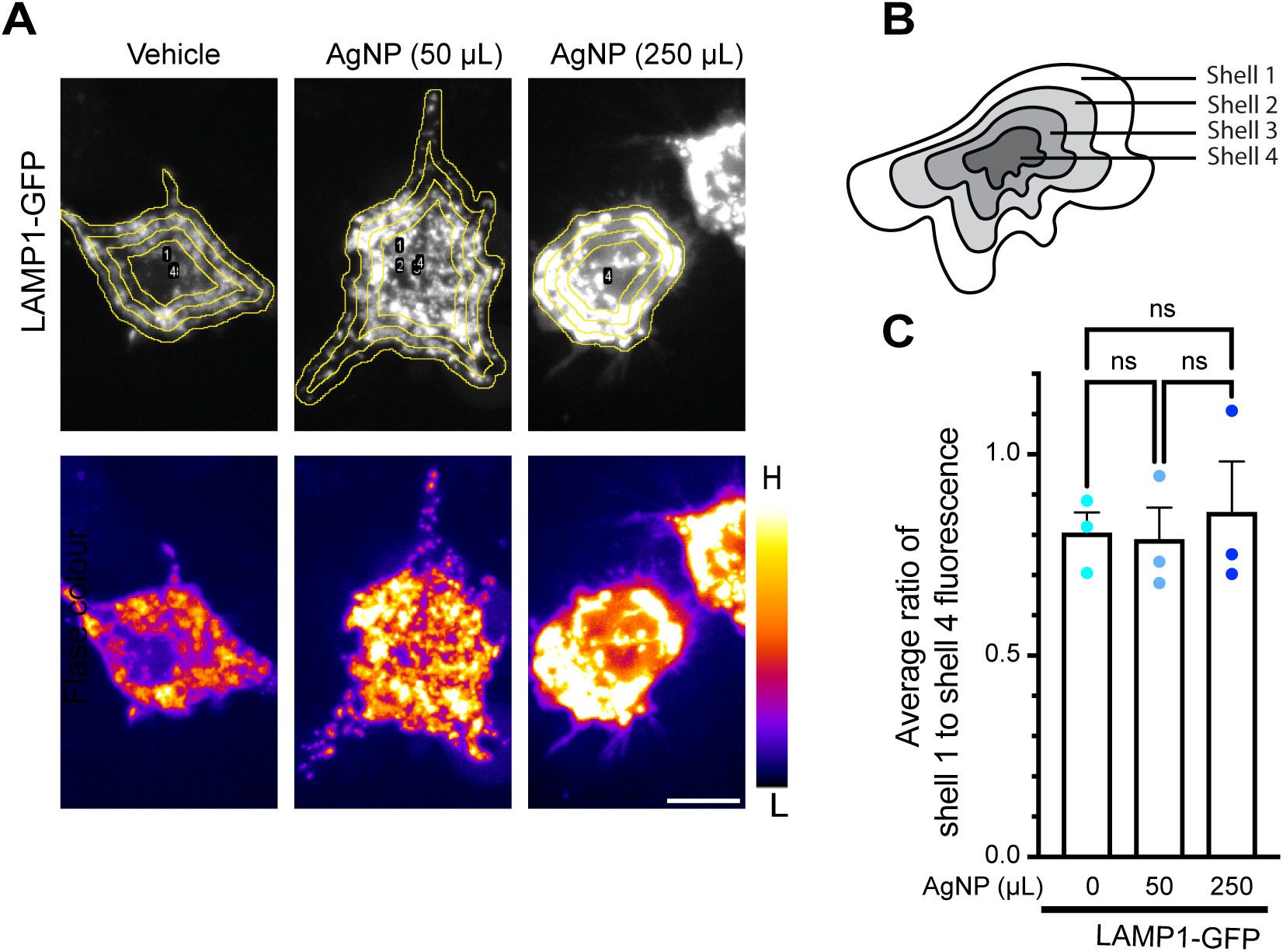
AgNPs do not affect lysosomal distribution in RAW macrophages. **A.** RAW macrophages were transfected with plasmid encoding LAMP1-GFP and exposed to vehicle, 50 µL AgNP, or 250 µL AgNP. Live-cells were then imaged by spinning disc confocal microscopy and are shown as grayscale (top row) or false-colour (bottom row) as before. Scale bar = 10 µm. B. Schematic of shell analysis showing four different regions of quantification. **C.** The mean ratio fluorescence between shell 1 to shell 4 was quantified and normalized to control cells with no AgNP. Shown is the mean ± SEM from N=3 independent experiments, where 20-30 cells per condition per experiment were quantified. Means were then tested by one-way ANOVA, followed by Tukey’s post-hoc test, where no significant difference was asserted.

Lastly, we examined lysosome distribution and remodelling in AgNP-treated RAW cells. First, using transient expression of LAMP1-GFP to demarcate lysosomes, we did not observe changes in lysosome distribution using a shell-analysis in cells loaded with AgNP relative to naive cells (Fig. 9). However, we observed that RAW cells loaded with AgNP appeared to have more basal tubular lysosomes labelled with fluorescent dextrans, though there was no inhibitory or additive effect when co-exposed with LPS to induce tubulation (Fig. 10; (Mrakovic *et al*., 2012)). Overall, under the observed conditions, AgNPs had little to modest effects on lysosome functions and properties.

**Figure 9:**
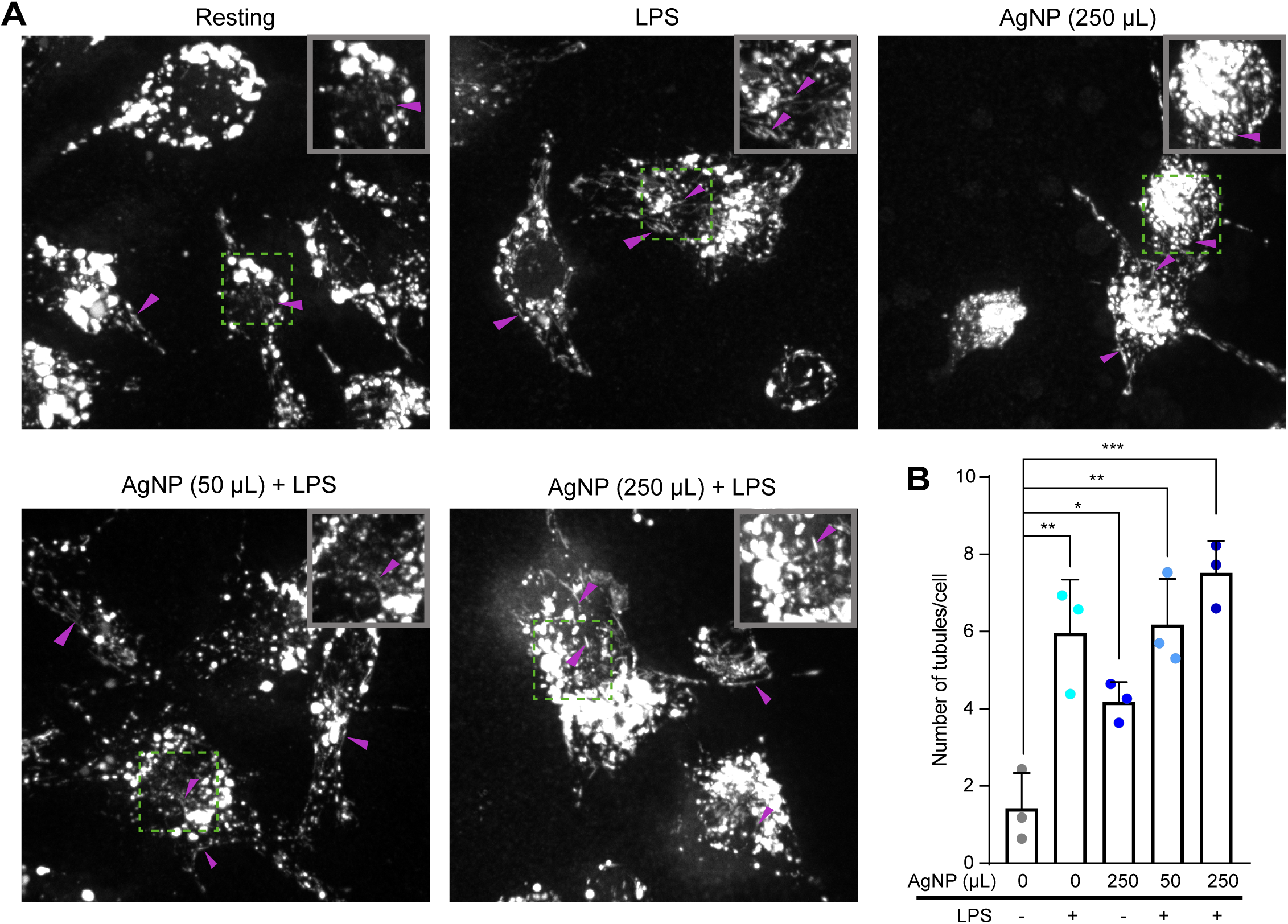
AgNPs induce lysosomal tubulation. **A**. Images of RAW macrophages were obtained through confocal spinning disk microscopy. Cells were labelled with 50 μg/mL of Alexa546- conjugated dextran for 1 h followed by a 1 h chase. Cells were then treated with LPS (500 ng/mL), or 250 μL of AgNP, or with 50 μL or 250 μL of AgNP for 1 h and then with LPS (500 ng/mL) for 2 h, or left untreated (Resting). Arrowheads point to tubules. Inset is about a 2x magnification. **B.** Mean number of lysosome tubules per cell. Each point represents the mean number of tubules from 25-30 cells in an independent experiment. The data was graphed as mean ± SEM. Statistical analysis was done using a one-way ANOVA, followed by a post-hoc Tukey’s comparison test; * indicates p<0.05, ** p<0.01, *** p<0.001.

**Figure 10:**
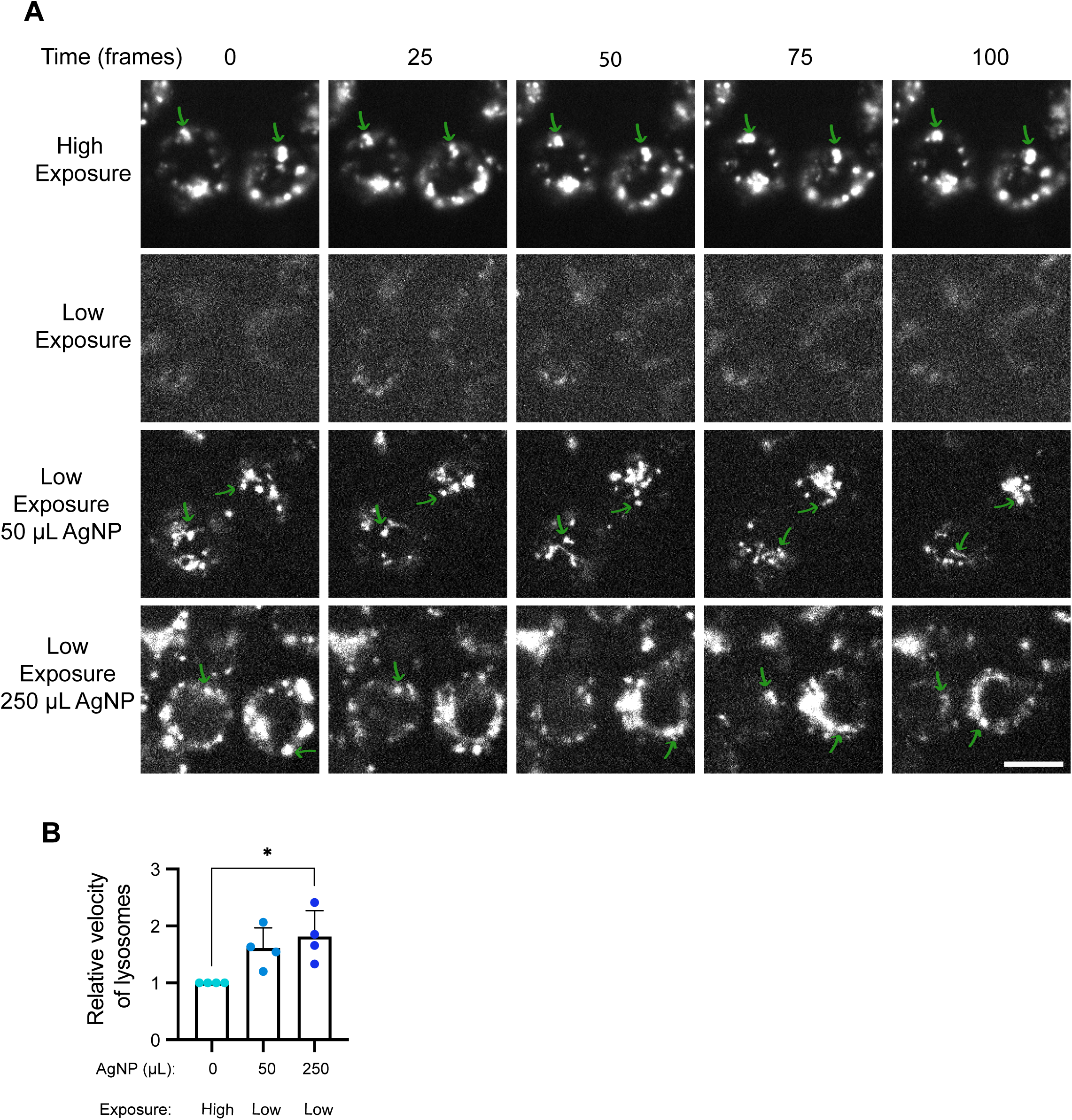
AgNP-enhanced fluorescence helps preserve lysosome motility during microscopy imaging. **A.** RAW cells were loaded with vehicle (top two rows) or AgNP (bottom rows) as before, followed by labelling with DQ-BSA. Cells were then imaged for 5 min at 0.2 frames/second at 10 mW (low) or 50 mW (high) laser power. Individual frames are shown at indicated times and correspond to Videos 5 through 8. Scale bar = 10 µm. Green arrows track individual lysosomes over time. **B.** Time sequences were subjected to particle tracking analysis to measure relative lysosome velocity using TrackMate. Velocity was normalized to control cells with no AgNP and high laser power. Shown is the mean velocity ± SEM from N=3 independent experiments, each with 5 cells per condition per experiment, or total of 15 cells. Means were tested by one-way ANOVA and Tukey’s post-hoc test. * indicates p<0.05.

### AgNP-enhanced fluorescence allows robust imaging conditions while protecting normal lysosome dynamics

We previously showed that photo-toxicity during spinning disc confocal imaging can interfere with lysosome dynamics due to ROS accumulation (Choy *et al*., 2018; Saffi *et al*., 2021). We hypothesized we could use AgNP-mediated fluorescence enhancement to lower photo-toxicity during imaging of lysosome dynamics in RAW cells. To test this, we labelled cells with DQ-BSA as before and tracked cells at high or low laser power over 5 min at 0.2 frames/s. We then subjected lysosomes to tracking analysis. Consistent with ROS-mediated damage that we observed previously (Choy *et al*., 2018; Saffi *et al*., 2021), while high laser power produces images with excellent signal-to-noise ratio, lysosomes became less motile in RAW macrophages (Fig. 11A, E, Video 1). In comparison, cells with low laser power displayed low signal-to-noise ratio and tracking lysosomes was not reliable (Fig. 11A, Video 2). However, cells that internalized 50 µL or 250 µL AgNP but were exposed with low laser power now displayed enhanced DQ-BSA signal-to-noise ratio (Fig. 11A, Videos 3 and 4). Importantly, we could track lysosome dynamics under these conditions, observing that lysosomes were significantly more motile and dynamic than in cells without AgNP particles and exposed to high light energy (Fig. 11B, Videos 1-4). Overall, we propose that AgNP-loading of lysosomes may be a useful tool to enhance fluorescence of lysosome-targeted probes, reducing the need for strong light energy or exposure, dropping photo-toxicity, which ultimately minimizes artifacts during live-cell imaging.

## Discussion

Fluorescence microscopy is a critical tool in biological studies and applications. However, it is hampered by the need to use excessive light energy to generate images with good signal-to-noise ratio. This ultimately leads to the generation of reactive oxygen species that can photobleach the fluorescent probes and cause be toxic to living samples. Photobleaching is not only inconvenient, but can limit observations, while phototoxicity can lead to artifactual observations (Tinevez *et al*., 2012; Waldchen *et al*., 2015; Boudreau *et al*., 2016; Han *et al*., 2017; Icha *et al*., 2017). Over the years, improved fluorochromes, camera and photodetector sensitivity, computer algorithms, and elegant design of dyes with suitable physicochemical properties have reduced the light intensity and/or exposure needed to obtain good signal-to-noise (Lidke and Lidke, 2012; Boutros *et al*., 2015; Jing *et al*., 2021). Despite these efforts, photobleaching and phototoxicity remain significant problems in live-cell imaging.

MEF holds great promise as a tool to augment spectroscopic and fluorescence-based applications (Lakowicz *et al*., 2003; Bouhelier *et al*., 2004; Ekgasit *et al*., 2004a; Kneipp *et al*., 2006a, 2006b; Petryayeva and Krull, 2011; Hodgson *et al*., 2020). Indeed, plasmonic metal nanoparticles can impact the behaviour of organic chromophores in several ways, including enhancing molecular fluorescence. From the perspective of the fluorochrome, this type of enhancement is originated by intensification of the incident electromagnetic field, via plasmonic excitation of the metallic nanostructures. Much of the relevant literature acknowledges the effects of nanoparticle shape, size, orientation, interparticle spacing or nanoparticle-dye separation upon ensemble averaged fluorescence enhancement factors (Lakowicz *et al*., 2002, 2008; Anger *et al*., 2006; Fort and Grésillon, 2008; Xie *et al*., 2008, 2013; Dragan and Geddes, 2011; Dutta Choudhury *et al*., 2012; Deng *et al*., 2013; Kinoshita *et al*., 2015; Lin and Chen, 2015; Fothergill *et al*., 2018; Wei *et al*., 2018). However, it is also critical to emphasize the importance of the optical properties of all the different components of the system i.e., fluorophores and nanoparticles working synergistically to achieve MEF. Interestingly, the spectral position of the plasmophore’s enhanced emission is nearly indistinguishable from that of fluorescence emission in free space: as such, applications of MEF abound for the magnification of fluorescence for bioimaging. For example, AgNP can enhance fluorescence of conjugated DNA oligomers (Lakowicz *et al*., 2003; Ekgasit *et al*., 2004a) and focal fields mapping (Bouhelier *et al*., 2004), while enhancement of single-molecule fluorescence by AgNP was exploited for the study of ribosomal mRNA translation (Bharill *et al*., 2011). Moreover, gold nanoparticles (AuNP) have been used to enhance single-molecule Raman spectroscopic imaging within cells, including within endosomes (Kneipp *et al*., 2006a, 2006b). However, to the best of our knowledge, AgNP have not been tested for their ability to enhance fluorescence intracellularly in living cells.

Herein, we showed that RAW macrophages fed AgNP by pinocytosis leads to enhanced fluorescence of BODIPY-conjugated lysosome-targeted probes, Alexa488-conjugated dextran, and GFP anchored to the cytoplasmic face of the lysosomal membrane as a LAMP1-GFP chimera. We demonstrate that this enhances signal-to-noise ratio affording a reduction in laser intensity, which abates phototoxicity, ultimately preserving cell function as seen in the preservation of lysosome motility. However, it should be noted that fluorescent probes differ in their properties such as quantum yield, molecular size, fluorochrome density, membrane association, and propensity for self-quenching. For example, Alexa488-dextrans are extended polysaccharides, while BODIPY-cholesterol likely preferentially associates with the membranes. These properties may affect how they interact with AgNPs, impacting the effectiveness of metal enhanced fluorescence. Thus, we recommend testing the doses of AgNPs and fluorochromes to optimize fluorescence enhancement for each fluorescent probe. Importantly, loading of AgNP within lysosomes had little effect on lysosome properties such as pH, degradative capacity, membrane integrity, autophagy, and distribution. We did observe an increase in lysosome tubulation in RAW macrophages, which could reflect a biological effect of AgNPs or that AgNPs contain impurities that may activate macrophages such as endotoxins. Thus, additional consideration is required if AgNPs were to be used to study lysosome tubulation.

Loading of AgNP into the endo-lysosomal system of cells is a promising tool to enhance fluorescence of endo-lysosomal probes, affording one to reduce light intensity and/or exposure. This is particularly important for imaging lysosome dynamics in macrophages, which have proven to be sensitive to photo-damage during spinning disc confocal microscopy and reactive oxygen species (Choy *et al*., 2018; Saffi *et al*., 2021). In fact, we have observed that lysosome motility and fusion-fission cycles can be altered during imaging and by reactive oxygen species (Choy *et al*., 2018; Saffi *et al*., 2021). Though light-induced alteration to lysosome function may depend on cell type, loading lysosomes with AgNP may thus be useful to visualize lysosome dynamics during confocal microscopy.

It is important to note again that the conditions for AgNP-mediated enhancing imaging of lysosomes will need optimization by the user, including consideration on the cell type under study, and the specific lysosomal probe. Moreover, future work is needed to explore the optimization of nanoparticles with different structure/properties relationships (e.g., size and shape) for which the plasmon resonance will need to be finetuned to enhance fluorescence of probes emitting at different wavelengths. Finally, AgNP may have different uses to enhance fluorescence in cells. Here, we used bulk pinocytosis of AgNP by macrophages to localize to lysosomes. However, it may be possible to enrich AgNP in endosomes with short pulse-chase periods. Alternatively, conjugating AgNP to ligands known to cycle through the endosomal system such as transferrin may allow for endosomal imaging (van Dam and Stoorvogel, 2002; Roberts *et al*., 2006). Similarly, AgNP may be conjugated to particulates like bacteria and polysterene beads to track phagosomes and study phagocytosis and processes like phagosome resolution (Fountain *et al*., 2021; Lancaster *et al*., 2021).

## Limitations of Study

Using our specific irradiation instruments, power, and ratio of reagents, we obtained highly reproducible AgNP suspension in terms of particle number, size, and shape. However, as these specifics may change between labs and users, initial production of AgNP by other workers will need to be tested and standardized. Moreover, while our study demonstrates that AgNP can enhance “green” fluorochromes within cells, future studies should develop AgNP with distinct spectral properties to determine if MEF can be achieved within cells when using fluorophores with different excitation/emission properties. Lastly, while loading AgNP into RAW macrophages did not impair cell growth or aggravate cell death within 24 h of observation, and several properties of lysosomes were not affected, we did observe an increase in basal tubulation in macrophages loaded with AgNPs. Thus, one should examine the impact of AgNP on specific functions of lysosomes or cell physiology processes to determine if their use is appropriate for that function.

## Acknowledgements

We acknowledge and thank Karol Golian and Lavinia Trifoi for assistance with nanoparticles synthesis. This work was supported by the Canada Research Chair Program (950-232333) and associated contributions from Toronto Metropolitan University, Project Grants from the Canadian Institutes of Health Research Project Grant (123373 and PJT-166047), the Canada Foundation for Innovation (32957) and associated contributions from the Ministry of Economic Development, Job Creation and Trade (32957) and Toronto Metropolitan University to R.J.B. In addition, this work was supported by Discovery Grants from the Natural Sciences and Engineering Research Council to C.N.A. (RGPIN-2016-04371) and S.I. (RGPIN-2018-04161) and the Toronto Metropolitan University Faculty of Science Dean’s Research Fund.

## Author Contributions

SAS is main contributor to this work carrying out most experimental design, data acquisition, interpretation, co-writing, and figure preparation.

AS, SH, NJ, and JC-P performed specific, targeted experiments, data acquisition, interpretation, and figure preparation.

GKH and MS established foundational data for this research and helped in experimental design. CNA, SI, and RJB are responsible for experimental design, resource acquisition, supervision, co-writing, figure preparation, and editing.

## Declaration of interests

Authors declare no conflicts of interest.

**Supplemental Video 1: Lysosome dynamics in RAW cells exposed to high laser power.** Cells without AgNP were labelled with DQ-BSA for 1 h. Cells were then imaged using 63x objective at 0.2 frames/second for 5 min using 50 mW laser power. Images had strong signal-to-noise ratio but lysosomes became less motile.

**Supplemental Video 2: Lysosome dynamics in RAW cells exposed to low laser power and without AgNP.** Cells without AgNP were labelled with DQ-BSA for 1 h. Cells were then imaged using 63x objective at 0.2 frames/second for 5 min using 10 mW laser power. Images had poor signal to noise ratio and lysosomes were difficult to track.

**Supplemental Video 3: Lysosome dynamics in RAW cells exposed to low laser power and 50 µL of AgNP.** Cells were loaded with 50 µL AgNP and labelled with DQ-BSA for 1 h. Cells were then imaged using 63x objective at 0.2 frames/second for 5 min using 10 mW laser power. Images had improved signal to noise ratio and lysosomes retained their dynamic behaviour.

**Supplemental Video 4: Lysosome dynamics in RAW cells exposed to low laser power and 250 µL of AgNP.** Cells were loaded with 250 µL AgNP and labelled with DQ-BSA for 1 h. Cells were then imaged using 63x objective at 0.2 frames/second for 5 min using 10 mW laser power. Images had improved signal to noise ratio and lysosomes retained their dynamic behaviour.

