## Supplemental Figure S1 for "Improved imaging and preservation of lysosome dynamics using silver nanoparticle-enhanced fluorescence"

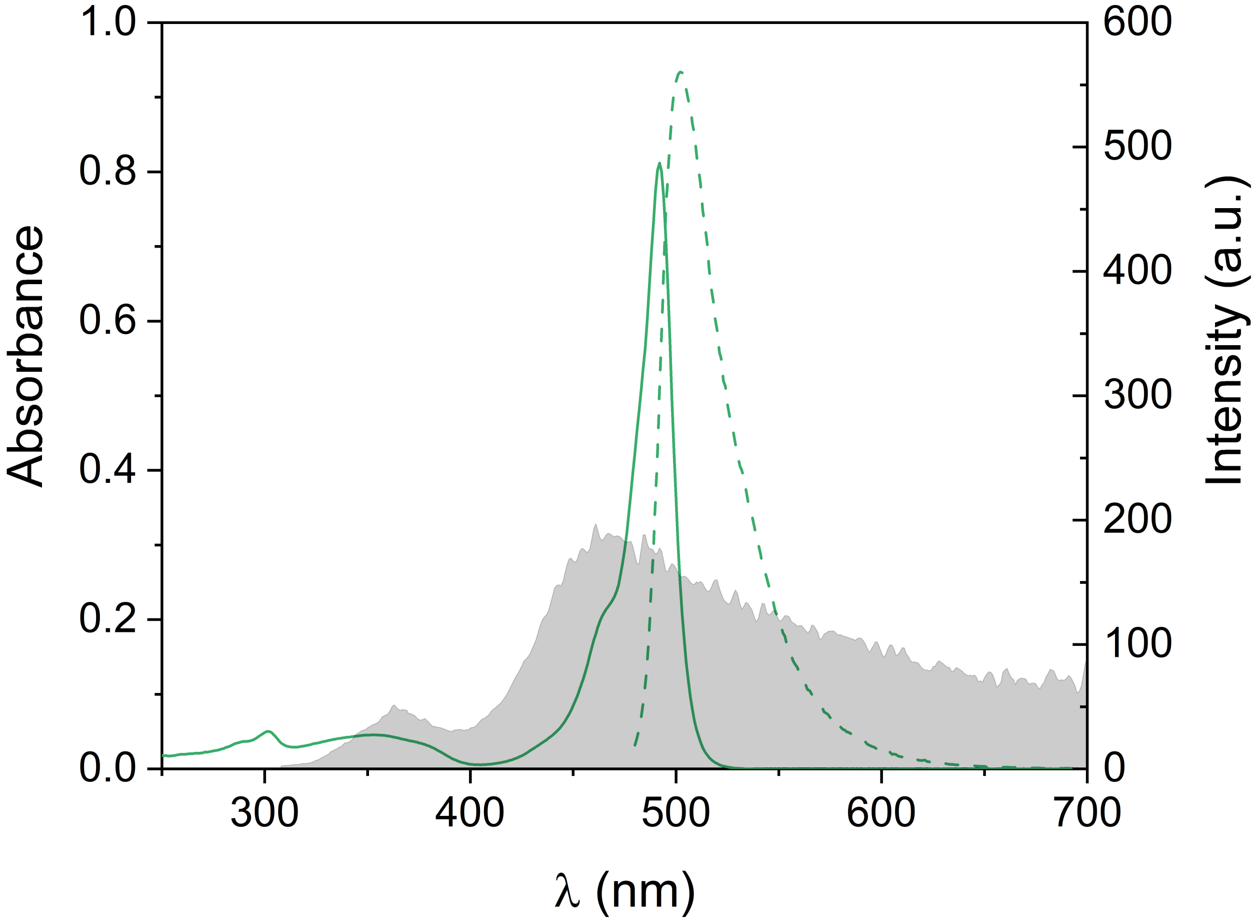


Figure S1. Absorption (solid green trace) and emission (dashed green trace) spectra (20 °C, CH3CN, λEx = 460 nm) of BODIPY493/503. Synchronous scattering (λEx = λEm, grey trace) spectrum of as-synthesized AgNP.
