## Supplementary figures and images for "Improved imaging and preservation of lysosome dynamics using silver nanoparticle-enhanced fluorescence"

### Supplemental Figure S2

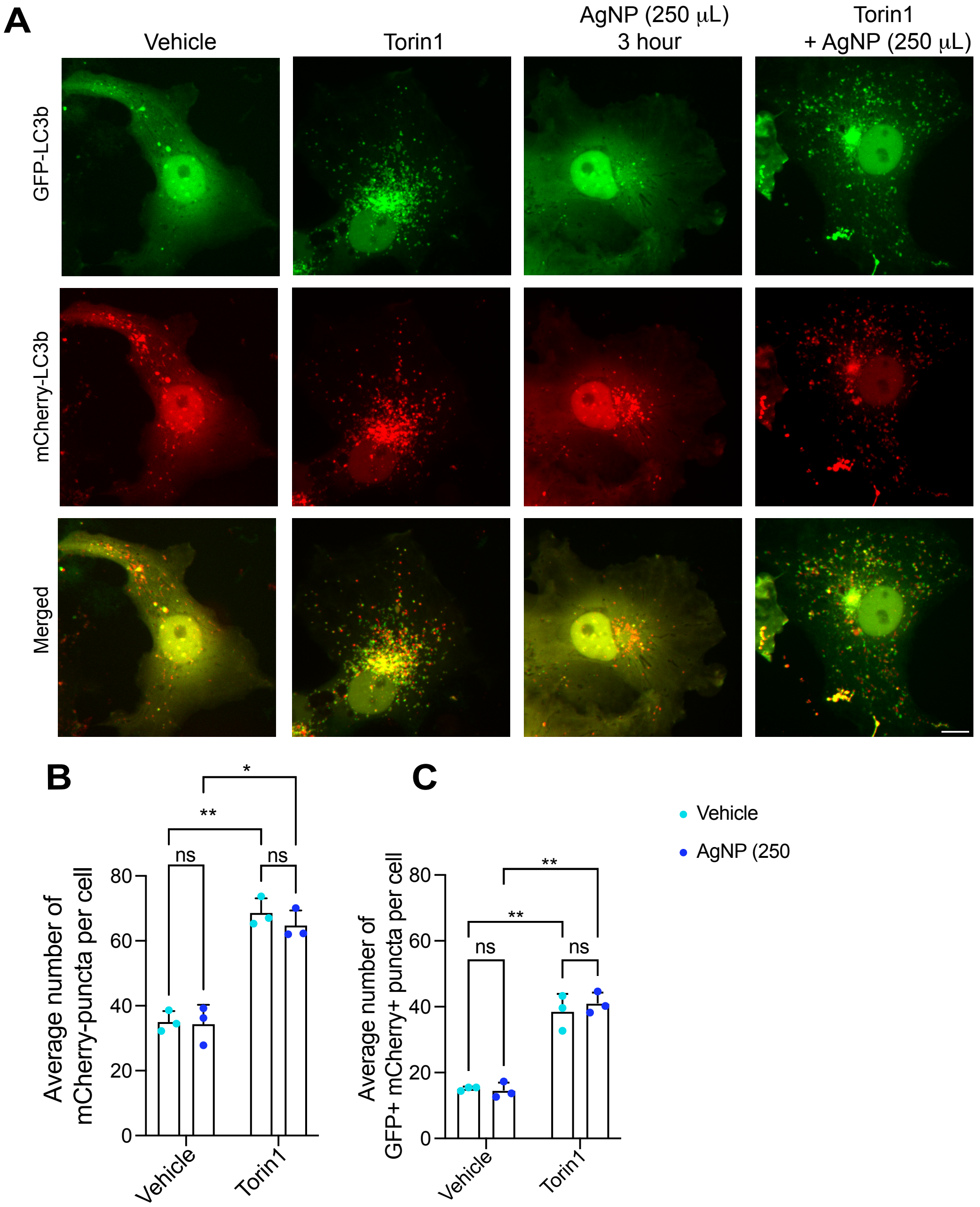
