## Supplemental Information for "Improved imaging and preservation of lysosome dynamics using silver nanoparticle-enhanced fluorescence"

**Supplemental Materials for Figures**

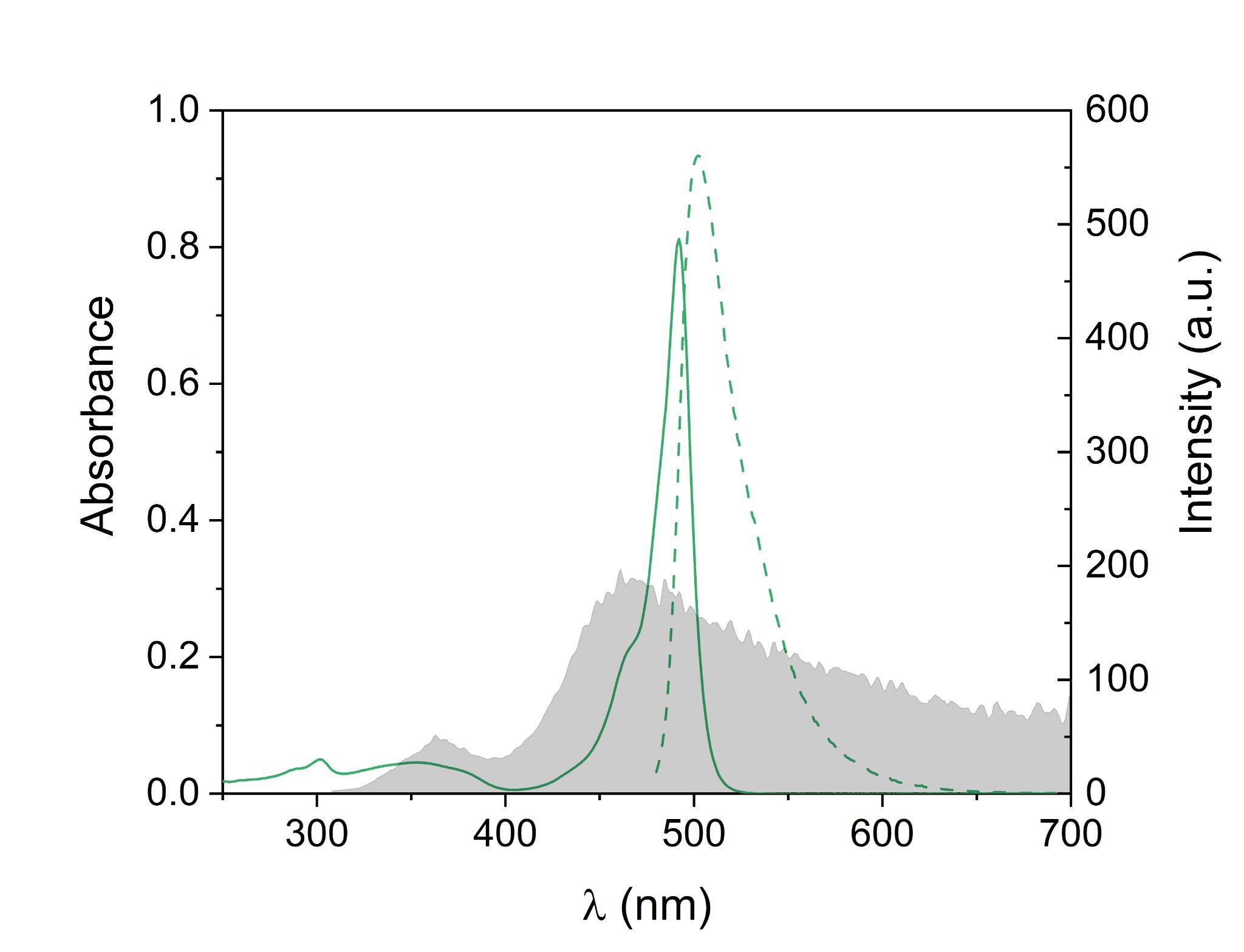

**Supplemental Figure S1.** Absorption (solid green trace) and emission (dashed green trace) spectra (20 °C, CH_3_CN, λ_Ex_ = 460 nm) of BODIPY^493/503^. Synchronous scattering (λ_Ex_ = λ_Em_, grey trace) spectrum of as-synthesized AgNP.

**Supplemental Figure S2: AgNP does not change basal autophagy or autophagic flux in COS7 cells.** COS7 cells were transfected with mCherry-GFP-LC3b and exposed to vehicle, 1 µM torin1, 250 µL AgNP, or 250 µL AgNP and 1 µM torin1 for 3 h, and then imaged live by spinning disc confocal microscopy. Scale bar = 10 µm. **A.** GFP-LC3b is shown in green (top), mCherry-LC3b in red (middle), and the superimposition of the two fluorophores in bottom) row. **B, C.** Mean number of mCherry puncta (B) and GFP+ mCherry+ puncta (C) were quantified from a minimum of 50 cells per condition per experiment. Shown is the mean ± SEM from N=6 independent experiments. One-way ANOVA and Tukey’s post-hoc test were used to compare means with no significant difference in autophagy and autophagic flux observed between.

**Supplemental Methods**

**Calculation of AgNP Concentration**

The concentration of silver nanoparticles (AgNP/mL) was estimated as follows. First, we calculated the weight of Ag (W_Ag_) used for nanoparticle synthesis. From a 0.8 mL of 10 mM AgNO_3_, and knowing the atomic weight of Ag = 107.87 a.u., we estimated W_Ag_ to be 8.63 x 10^-4^ g. As the density of silver is d_Ag_ = 10.49 g/cm^3^, the volume of Ag used is:

V_Ag_ = (8.63 x 10^-4^ g)/(10.49 g cm^-3^) = 8.227 x 10^-5^ cm^3^

According to TEM measurements, the average size of our AgNP is 2*r* = 3 nm, where *r* is the nanoparticle radius.^S1^ As such, the volume of a single spherical AgNP is:

V_AgNP_ = 4/3 π *r*^3^ = 4/3 π (1.5 nm)^3^ = 14.13 nm^3^

We can then calculate the total number of AgNP (N_AgNP_) in the colloidal solution (40 mL) as:

N_AgNP_ = V_Ag_/V_AgNP_ = (8.227 x 10^16^ nm^3^)/(14.13 nm^3^) = 5.822 x 10^15^

and the concentration of AgNP/mL as:

(5.82 x 10^15^)/40 mL = **1.456 x 10^14^ AgNP/mL**

We validated these calculations using a second independent method*.*^S2^ The amount of AgNP formed when 0.8 mL of 10 mM AgNO_3_ is reduced can be calculated as:

moles of AgNO_3_ = (0.8 x 10^-3^ L)(10 x 10^-3^ mol L^-1^) = 8 x 10^-6^ mol

No. of Ag atoms (N_atom_) = (8 x 10^-6^ mol)N_A_ = (8 x 10^-6^ mol)(6.022 x 10^23^ mol^-1^) = 4.818 x 10^18^

where *N_A_* is the Avogadro constant. The number of AgNP can then be assessed as:

N_AgNP_ = N_atom_/N

where *N* is the number of silver atoms per nanoparticles, and which is equal to:

N = (*r*_AgNP_/*r*_atom_)^3^ = (1.5 nm/0.144 nm)^3^ = 1130

Thus,

N_AgNP_ = N_atom_/N = (4.818 x 10^18^)/1130 = 4.263 x 10^15^

and,

(4.263 x 10^15^)/40 mL = **1.066 x 10^14^ AgNP/mL**

Which is in excellent agreement with the estimated value above. Averaging the results obtained with the two independent methods, we can conclude that concentration of AgNP is reasonably estimated as **1.261 x 10^14^ AgNP/mL**.
